# Noise properties of adaptation-conferring biochemical control modules

**DOI:** 10.1101/2023.02.05.525388

**Authors:** Brayden Kell, Ryan Ripsman, Andreas Hilfinger

## Abstract

A key goal of synthetic biology is to establish functional biochemical modules with network-independent properties. Antithetic integral feedback (AIF) is a recently developed control module in which two control species perfectly annihilate each other’s biological activity. The AIF module confers robust perfect adaptation to the steady-state average level of a controlled intracellular component when subjected to sustained perturbations. Recent work has suggested that such robustness comes at the unavoidable price of increased stochastic fluctuations around average levels. We present theoretical results that support and quantify this trade-off for the commonly analyzed AIF variant in the idealized limit with perfect annihilation. However, we also show that this trade-off is a singular limit of the control module: Even minute deviations from perfect adaptation allow systems to achieve effective noise suppression as long as cells can pay the corresponding energetic cost. We further show that a variant of the AIF control module can achieve significant noise suppression even in the idealized limit with perfect adaptation. This atypical configuration may thus be preferable in synthetic biology applications.

---

The behaviour of even the most complex electrical circuit can be perfectly predicted because components such as resistors, inductors, and capacitors have network independent properties. In contrast, the behavior of complex chemical reaction networks in cells is extremely challenging to analyze due to the lack of well-defined modules. Identifying biochemical modules with network independent properties remains one of the key challenges for synthetic biology. One biochemical module that has received much recent theoretical and experimental attention [1–26] is the idealized antithetic integral feedback (AIF) control module in which a sensor species and a reference species perfectly annihilate each other’s biological activity [1, 8]. This module confers robust perfect adaptation [1, 8, 27] to any intracellular component of interest regardless of that component’s uncontrolled dynamics: Any intracellular component of interest controlled by the AIF module will perfectly readjust its steady-state average in the face of sustained perturbations as long as the overall system is stable [1, 8, 27].

In addition to robustness that guarantees constant average abundances in different environments, cells must also manage stochastic fluctuations around averages in a given environment [28–31]. A priori, the former kind of robustness says nothing about the latter. However, recent work has suggested that the robust perfect adaptation of averages conferred by the AIF module necessarily increases stochastic noise around averages [2, 32, 33].

Here, we present theoretical results that support and quantify this trade-off for the commonly analyzed idealized variant of the AIF control module. However, we further show that this behaviour is a singular limit of the idealized control module that does not describe the noise properties of realistic control in cells where the AIF control molecules necessarily undergo (some) individual degradation [4, 6, 17].

Our results suggest that the non-idealized AIF module can suppress noise significantly below open-loop levels while simultaneously conferring *near-*perfect adaptation of the average. We further show that an atypical variant of the AIF control module can achieve significant noise suppression even in the idealized limit with perfect adaptation. This suggests that not all AIF topologies are equal and that certain implementations of AIF control are preferable in synthetic biology applications due to their desirable noise properties.

While our results highlight the lack of a fundamental trade-off between the robustness of average abundances and stochastic fluctuations around averages, they also establish the large energetic burden required to simultaneously reduce noise and sensitivity. This burden may place practical limitations on how small these quantities can be made experimentally. However, if both small noise and small sensitivity are required, our results indicate that the remarkable AIF module conceived by Briat, *et al*. [1] can – in principle – get the job done without the need for additional feedback control.

## RESULTS

### Unavoidable noise penalty of idealized reference-actuated antithetic integral feedback

Achieving robust perfect adaptation for ensemble averages in discrete stochastic processes requires components whose average elimination fluxes are perfectly matched at stationarity [8]. In biochemical systems, this condition can be realized through two control molecules that annihilate each other’s biological activity through complex formation [1, 8] (Fig. 1).

**Figure 1.**
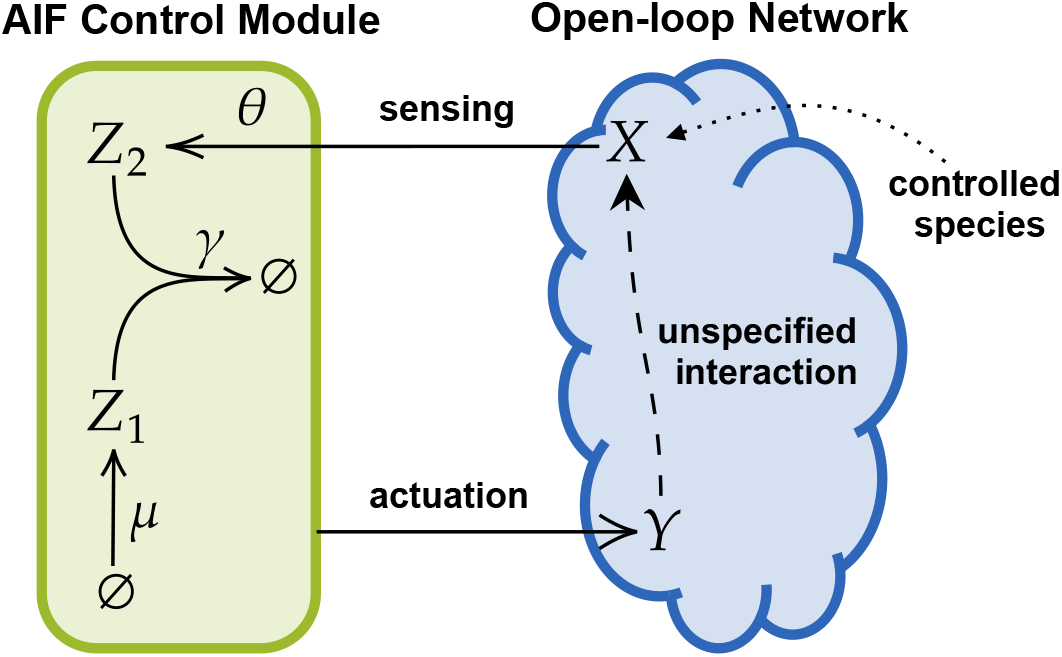
Minimal antithetic integral feedback (AIF) control module. The AIF module (green) utilizes two control species to confer robust perfect adaptation to a species of interest *X* within an unspecified biochemical network (blue): A reference species *Z*_1_ is produced at a fixed rate *μ* and a sensor species *Z*_2_ is produced at a rate *θx* proportional to the abundance of the controlled species. The two control species react irreversibly to form an inert complex. The feedback loop is closed by an actuation reaction that directly or indirectly affects *X*-levels and depends on *Z*_1_ or *Z*_2_ abundances. As long as the closed-loop dynamics are stable, the steady state average abundance of *X* is set by the control module and independent of any reaction rates of the controlled network [1]. AIF control thus rejects sustained perturbations to a controlled component regardless of its uncontrolled dynamics as long as perturbations do not affect overall stability or the set-point encoded by the ratio *μ/θ* (see Eq. 3).

Such an antithetic integral feedback (AIF) control module includes a reference species *Z*_1_ produced at a fixed constant rate and a sensor species *Z*_2_ that is produced at a rate proportional to the abundance of the controlled species *X*. The minimal idealized AIF control module [1] is thus given by the following stochastic transitions of the control species:

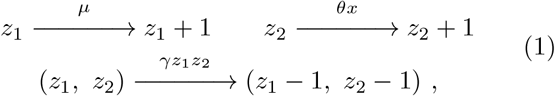

where *x, z*_1_, *z*_2_ denote the instantaneous abundances of the respective species *X, Z*_1_, *Z*_2_. The feedback loop is closed by an (unspecified) actuation reaction that directly or indirectly affects *X*-levels and depends on *Z*_1_ or *Z*_2_ abundances (Fig. 1).

Within the AIF module, the ensemble average dynamics of the control species follow

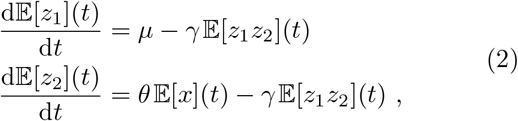

where 𝔼[·] denotes the expectation operator over the joint probability distribution of *x, z*_1_, *z*_2_ across an ensemble of isogenic cells in a shared environment. If the closed-loop dynamics are stable, the stationary state condition implies that

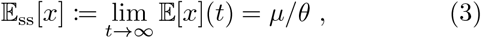

regardless of the dynamics or parameters of additional processes affecting *X*-levels. The stationary ensemble average of *X* thus perfectly tracks the set-point *μ/θ* regardless of other processes affecting *X*, as long as perturbations do not affect the ratio *μ/θ* and closed-loop stability is maintained. This remarkable property is referred to as robust perfect adaptation, since perfect set-point tracking at stationarity is robust to unknown details or sustained perturbations to the controlled network under stable conditions [1, 8, 27].

However, anecdotal numerical data for specific realizations of reference-actuated AIF systems [1, 2] and analytical results in the *γ* ≫ *μ* limit [2] suggest that reference-actuated AIF control leads to increased stochastic fluctuations of *X*-levels around their average compared to an open-loop control strategy maintaining the same average *X*-level. To investigate the noise properties of AIF control, we first consider the following minimal reference-actuated system where molecules of the species of interest *X* undergo the following production and degradation reactions:

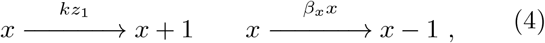

where the reactions within the control module are given by Eq. 1 (see Fig. 2A).

**Figure 2.**
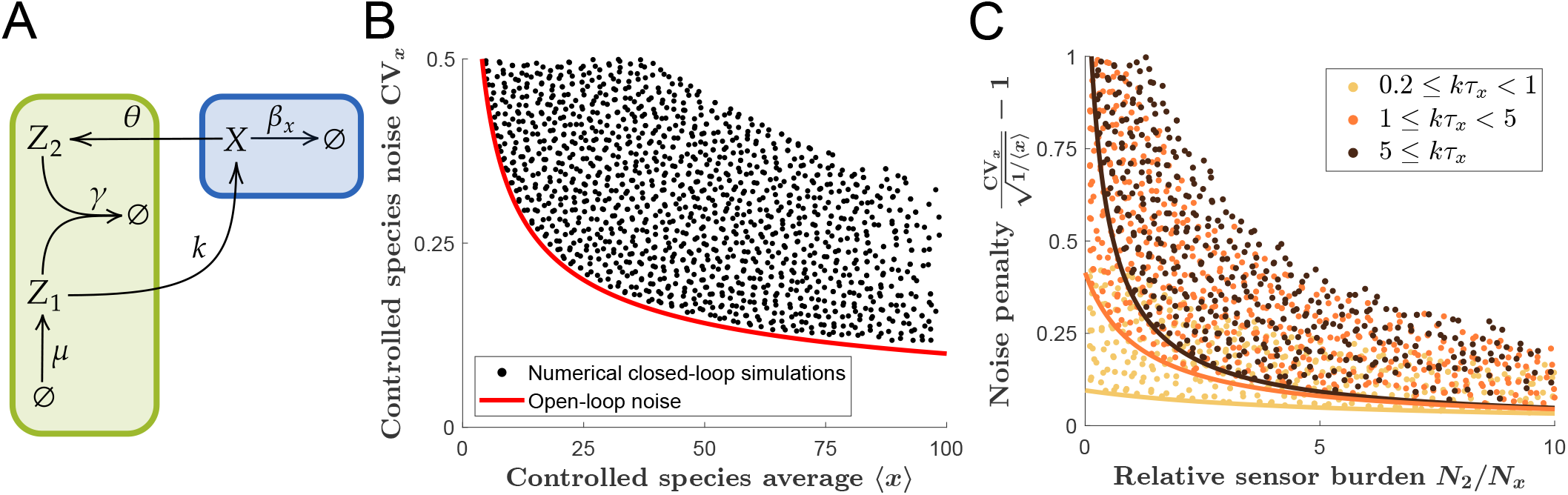
Idealized reference-actuated antithetic integral feedback (AIF) confers an unavoidable noise penalty. A) We consider a minimal model of idealized reference-actuated AIF control where the reference species *Z*_1_ influences the controlled species *X* through a production rate proportional to the abundance of *Z*_1_ molecules, as defined by Eqs. 1 and 4. Although the controlled species *X* undergoes first-order degradation its average abundance perfectly tracks the AIF-conferred set-point *μ/θ* regardless of the value of its degradation rate. B) Eq. 6 suggests that the controlled species noise must exceed its open-loop noise given by Poisson fluctuations (red line). A systematic numerical exploration involving over 10^6^ parameter sets spanning four orders of magnitude (see SI) confirmed this analytical prediction. No violations were found and black dots correspond to a representative sub-sample of our numerical simulations to indicate the accessible space. C) The minimum noise penalty depends on how many controller molecules are made per controlled molecule on average, given by (*N*_1_ + *N*_2_)*/N*_*x*_ = 2*N*_2_*/N*_*x*_, where *N*_1_, *N*_2_, and *N*_*x*_ denote the average number of birth events of *Z*_1_, *Z*_2_ and *X* molecules during the average lifetime of the controlled species. The minimum noise penalty of Eq. 6 is shown for three different values of the actuation gain *k* (solid lines). Numerical simulations (coloured dots) show that for each value of *k* the inequality based on linear approximations correctly bounds the behavior of all systems in the regime ⟨ *x* ⟩ ≥ 1. Shown is a representative sub-sample of the all simulated points with ⟨ *x* ⟩ ≥ 1 that indicates the respective accessible space. See SI for an evaluation of numerical sampling error verifying the lack of violations of the predicted bounds.

Previous work [1] has established the ergodicity of the above system such that

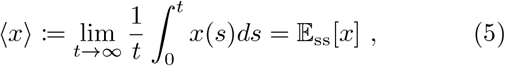

where ⟨*x*⟩ denotes the stationary single-cell time average and *x*(*t*) denotes the time-varying abundance of *X* in a single cell.

In this minimal model the species of interest *X* ex-hibits Poisson fluctuations in the absence of AIF con-trol. Its open-loop coefficient of variation is thus given by 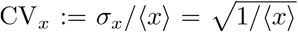, where *σ*_*x*_ is the standard deviation of *X*-levels at stationarity. To quantify the stationary fluctuations of *X*-levels under AIF control we utilize the standard linear noise approximation [34–37] to approximate the stationary covariance matrix for this system (see Materials and Methods). Under this approx-imation, we find (see SI) that all possible solutions satisfy the inequality

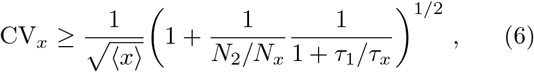

where *N*_*x*_ and *N*_2_ denote the average number of birth events of *X* and *Z*_2_ molecules, respectively, during the average lifetime *τ*_*x*_ = 1*/β*_*x*_ of the controlled species, and *τ*_1_ denotes the average lifetime of *Z*_1_ molecules.

Eq. 6 predicts that the noise in the controlled species levels is always larger under AIF control compared to the open-loop fluctuations. However, due to the non-linearity of the complex formation reaction rate in the AIF module (see Eq. 1), the linear noise approach is only approximate and not guaranteed to describe systems beyond the infinitesimal noise limit. We thus utilize Gillespie’s exact stochastic simulation algorithm [38, 39] to numerically determine the stationary state fluctuations of the above system with arbitrary precision (see Materials and Methods). Simulation results from a systematic parameter search with more than 10^6^ parameter sets spanning four orders of magnitude of separation between individual parameter values (see SI) suggest the AIF-controlled systems indeed exhibit higher noise levels than their open-loop counterparts in all noise regimes (Fig. 2B).

Furthermore, Eq. 6 quantifies how the minimum noise penalty depends on the details of the AIF control. This noise penalty can be reduced by increasing either *N*_2_*/N*_*x*_ or *τ*_1_*/τ*_*x*_. For a fixed set-point, increasing the former corresponds to increasing the average production flux of control species molecules, which is directly proportional to the overall energetic cost of the control module [1]. In practice, increasing *τ*_1_*/τ*_*x*_ is achieved by increasing *τ*_1_ since *τ*_*x*_ is an intrinsic parameter of the controlled process. For for a given *N*_2_*/N*_*x*_, increasing *τ*_1_ requires decreasing the actuation rate constant *k*. Previous work has suggested that the adaptation settling time diverges when this rate constant becomes small [10]. Eq. 6 thus predicts an energy-noise trade-off for this system that can only be overcome by infinitely slow adaptation times, which is borne out by our numerical simulations, as long as ⟨*x*⟩≥ 1 (Fig. 2C). See SI for details of the numerical approach to rule out violators for ⟨*x*⟩≥ 1.

An energy-speed-noise trade-off similar to the trade-off described by Eq. 6 has been discussed in the context of a mean-field model of sensory adaptation with Langevin dynamics, where fluctuations are modelled as extrinsic white noise [40]. While such mean-field approaches have been useful to understand sensory adaptation in, for example, bacterial chemotaxis [40–42], they do not describe the trade-off of Eq. 6 because their noise strength is an extrinsic model parameter rather than an outcome of the control process itself. Further note that in the presence of intrinsic noise, adaptation in bacterial chemotaxis is at best near-perfect [43], whereas robust perfect adaptation is a structural property of AIF control regardless of intrinsic noise levels [1, 8, 33].

Recent theoretical and computational work has established that adding supplemental control structures to the above reference-actuated AIF module can reduce stochastic noise compared to AIF control without the additional control [2, 9]. However, these results do not negate the existence of the above noise penalty due to AIF control because they do not compare noise levels of AIF controlled systems to their uncontrolled levels.

### Reference-actuated antithetic integral feedback can achieve near-perfect adaptation and noise suppression

In the previous section we established how an increase in stochastic noise is a necessary side effect of achieving *perfect* adaptation through the idealized reference-actuated AIF control module. However, real implementations of AIF can never confer perfect adaptation because intracellular components in growing and dividing cells are inevitably subject to dilution [4, 6, 17].

Under the assumption of exponentially growing cells, the dilution of intracellular components can be modelled as a first-order decay process where the rate constant is given by the specific growth rate of the host cell [6, 17, 44–46]. Additionally, the controller species of the AIF module may be subject to degradation. Thus, we next consider the minimal reference-actuated AIF topology described by Eqs. 1 and 4 with the following additional first order decay reactions:

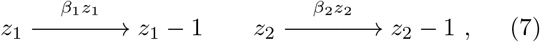

where *β*_1_, *β*_2_ are effective decay rate constants due to dilution and degradation of *Z*_1_, *Z*_2_ molecules (Fig. 3A). These decay processes result in ‘leaky’ integration and imperfect adaptation [4, 6, 17]. Prior work established that in the limit of *μ, θ, γ* ≫ *β*_1_, *β*_2_, the degradation fluxes of Eq. 7 are negligible and the set-point tracking error can be made arbitrarily small [6]. However, biochemical and energetic constraints limit how large controller reaction rates can be in a given experiment. Furthermore, in practice the value of *μ* and *θ* may not be known, which makes the set-point tracking error experimentally unobservable. Prior experimental work thus did not measure the set-point tracking error but focused on verifying the robustness conferred by the AIF control module by quantifying the sensitivity of average *X*-levels to perturbations in the degradation rate of that species [8]. Mirroring this experimental approach we consider the sensitivity of the stationary average ⟨*x*⟩ to perturbations in the controlled system parameter *β*_*x*_, defined as

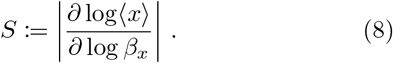

**Figure 3.**
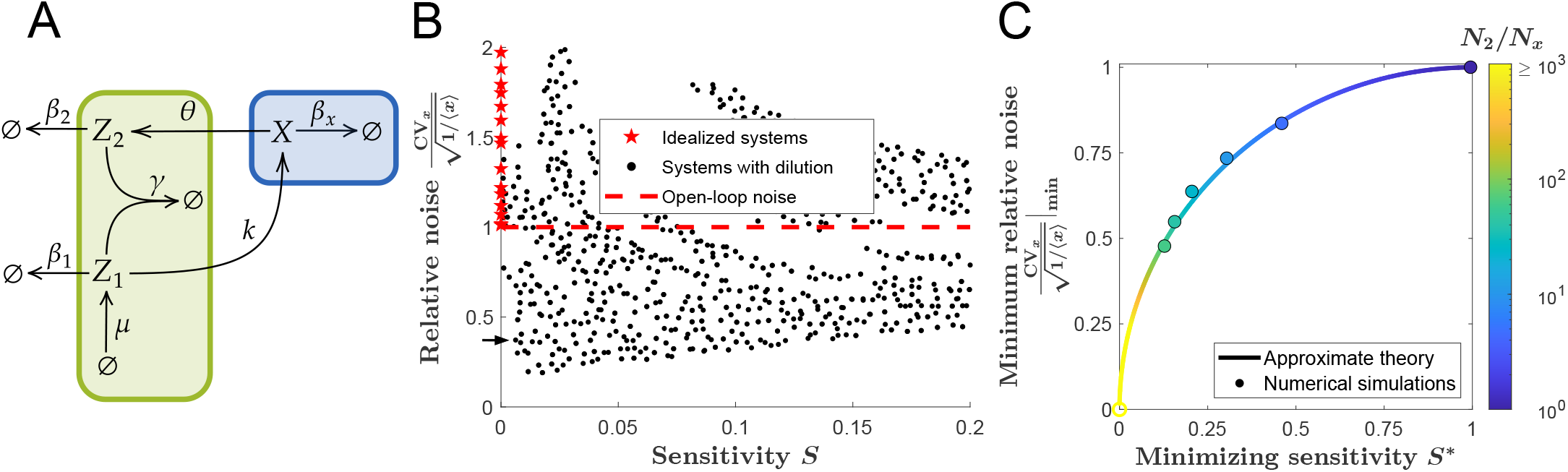
Non-idealized reference-actuated antithetic integral feedback control can simultaneously suppress noise and confer near-perfect adaptation. A) We consider the same reference-actuated AIF control system as in Fig. 2 but now subject control molecules to first-order degradation (Eq. 7) to represent the inevitable degradation and dilution effects in cells. All such non-idealized systems exhibit a non-zero sensitivity of the average abundance of the controlled species *X* to changes in its degradation rate. B) Shown are noise levels and sensitivity coefficients *S* as defined in Eq. 8 for a sub-sample of numerical simulations with different control parameters (see Materials and Methods; see also SI). Non-idealized systems (black dots) can achieve significant noise suppression levels for even minute deviations from perfect adaptation. For example, numerically we found one control system (indicated by arrow) that achieved a sensitivity of *S* ≈ 0.0062 while suppressing noise to 37% of the open-loop levels (*μ* = 442, 000, *θ* = 10, 000, *k* = 1.65, *γ* = 165, 000, *β*_*x*_ = 1, *β*_1_ = *β*_2_ = 10). In contrast, the previously considered (Fig. 2B) idealized control systems (red stars) always exhibit larger noise than the open-loop system (dashed red line). C) For a given minimum control burden *N*_2_*/N*_*x*_, the minimum noise level is predicted to be achievable only for a particular sensitivity *S*\* (see SI). Shown are the minimum CV_*x*_ and corresponding *S*\* values observed in numerical simulations with *N*_2_*/N*_*x*_ = 1, 5, 10, 20, 40, 80. The points are coloured according to the prediction *N*_2_*/N*_*x*_ = 1*/*(*S*\*)^2^ using the numerical *S*\* values, showing close agreement with the relationship predicted by Eq. 10. Numerically, we find *N*_1_ ≫ 1 is required for CV_*x*_ to approach the predicted minimum, where *N*_1_ is the average number of *Z*_1_ birth events during the average lifetime of the controlled species. We further observed that the value of *N*_1_ required to approach the minimum increased with *N*_2_*/N*_*x*_ (see SI), which made simulations too computationally intensive for an exhaustive parameter search to find the theoretically achievable minima for a given *N*_2_*/N*_*x*_ in the regime *N*_2_*/N*_*x*_ *>* 80.

Perfectly adapting systems satisfy *S* = 0, but any physi-cal realization of AIF will exhibit *S >* 0 due to dilution and degradation of control species molecules.

Sensitivity coefficients can be readily calculated for steady-states of deterministic systems. However, in stochastic systems the non-linear complex formation rate of the AIF module (see Eq. 1) results in a lack of moment closure, precluding exact analytical calculation of the stationary sensitivity coefficient (see SI). Following a finite difference approach [47, 48], we thus determined *S* numerically for a given system by measuring the effect of 1% perturbations in *β*_*x*_ on the numerical simulation average ⟨*x*⟩ (see Materials and Methods).

Through a numerical exploration of systems with ran-domized parameters (see SI) we find that systems with near-perfect adaptation have different noise properties from the idealized limit (Fig. 3B). For example, a system with a minuscule sensitivity of *S* ≈ 0.0062 can already achieve a noise reduction of over 60% relative to the uncontrolled system. The noise penalty quantified in the previous section, and declared a general feature of AIF control [2, 32, 33], thus seems a singular feature of the non-physical idealized case that does not characterize systems with small but non-zero sensitivities (Fig. 3B). Inevitable numerical sampling error can significantly affect numerical sensitivity values in the near-perfect adaptation regime. We thus plot the average sensitivity and noise values obtained when each parameter set was independently simulated three times to obtain reliable estimates. See SI for an evaluation of the sampling error of these data that confirms the above conclusions.

Analytically determining the sensitivity *S* exactly is impossible for the above systems. However, approximat-ing the stationary covariance between *Z*_1_-and *Z*_2_-levels using the linear noise approximation leads to a closed system of equations for the stationary state fluxes which in turn can be used to derive an approximation for *S* (see SI). Additionally, the linear noise approximation predicts (see SI) that avoiding a noise penalty requires *N*_2_/*N*_*x*_ ≥ 1, for which noise suppression is bounded by

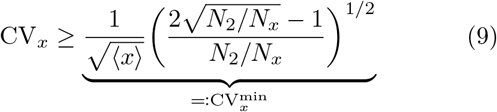

A systematic numerical exploration of systems with random parameters (see SI) suggests that the predic-tion of Eq. 9 based on the linear noise approximation constrains all possible systems (see SI). The numerical data additionally suggest that for a fixed value of *N*_2_*/N*_*x*_ there is a trade-off between sensitivity and noise un-til the noise floor 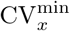 is achieved for some value of *S* = *S*\* (see SI). Our analytical approximations predict 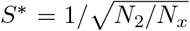 (see SI), which leads to the following relation between the noise floor and *S*\*:

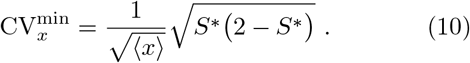

Reaching the predicted noise floor becomes exceedingly expensive computationally as *N*_2_*/N*_*x*_ increases, and thus we do not systematically explore the entire accessible space for *N*_2_*/N*_*x*_ > 80 (see SI). Numerically, we were thus unable to enter the corner in Fig. 3B of infinitesimally small noise and sensitivity. However, in the parameter regime *N*_2_*/N*_*x*_≤80 in which we are confident that we numerically sampled the entire accessible space (see SI) our exact numerical simulations indeed follow Eq. 10 (Fig. 3C). It is thus tempting to extrapolate the validity of Eq. 10 all the way to the corner of infinitesimal noise and sensitivity in the limit of *N*_2_≫*N*_*x*_. How-ever, this extrapolation into the corner of Fig. 3B remains not strictly proven and the numerical data indicated in Fig. 3B are the closest systems we have observed. While our results suggest that there is no fundamental limit on noise suppression for a given sensitivity, they also establish that achieving large noise suppression for small sensitivities requires an extremely large production of control molecules because the minimum noise level decreases only with the quartic root of *N*_2_*/N*_*x*_. The energetic bur-den and potential resource competition due to making a large number of control molecules, might negatively affect organismal fitness [49, 50] and diminish the performance of synthetic circuits [51–53] unless control molecules are very small and cheaply made.

Note, the addition of the dilution reactions of Eq. 7 results in unequal average production fluxes for the control species such that *N*_2_ no longer uniquely de-termines the average total production flux of control species. However, the flux of *Z*_2_ molecules still defines a *minimum* average production flux of control species.

### Sensor-actuated antithetic integral feedback can suppress noise in both the idealized and non-idealized case

While much of the literature on AIF control has focused on reference-actuated systems, different actuation inputs are possible [8, 9, 23, 54, 55]. In particular, the antithetic integral ‘rein controller’ proposed in [9] considers AIF control where both the reference species *Z*_1_ and sensor species *Z*_2_ affect the controlled species *X*.

To contrast the noise properties of different AIF variants, we consider the AIF control module of Eq. 1 where *only* the sensor species *Z*_2_, instead of the reference species *Z*_1_, affects the the controlled species of interest *X* (Fig. 4A), i.e., we replace the previously considered dynamics of *X* defined in Eq. 4 by

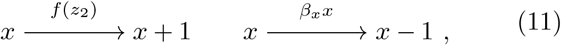

where *f* is a decreasing function of *z*_2_. The previous closed-loop ergodicity theorem strictly applies only to reference-actuated AIF control [1]. However, we find for all numerical simulations of idealized sensor-actuated systems that the time-average of individual time-traces agrees with the ensemble average *μ/θ* guaranteed by perfect adaptation, suggesting closed-loop ergodicity (see SI).

**Figure 4.**
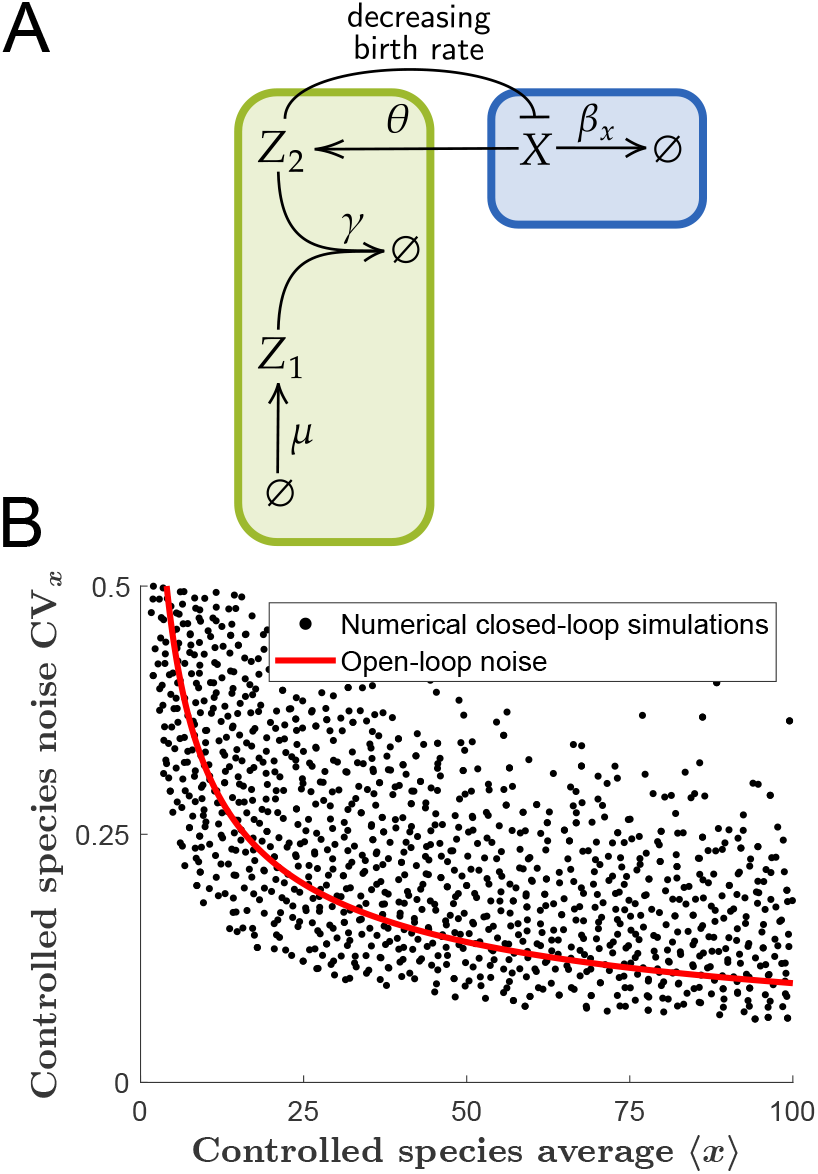
Sensor-actuated antithetic integral feedback (AIF) can simultaneously suppress noise below open-loop levels while conferring perfect adaptation. A) We consider a minimal model of idealized sensor-actuated AIF control where the sensor species *Z*_2_ suppresses the pro-duction of the controlled species *X* (see Eq. 11). The controlled species undergoes first-order degradation and its averfectly tracks the set-point *μ/θ* regardless of the value of its degradation rate *β*_*x*_. B) Eq. 12 suggests that sensor-actuated AIF controllers can suppress noise levels of the controlled species below their open-loop levels given by Poisson fluctuations (red line). Numerical simulations (black dots) confirm this analytical prediction. Shown is a representative sub-sample of sensor-actuated AIF controllers. Note, that all simulations achieve perfect adaptation, and the apparent envelope of the data does not correspond to a fundamental limit but is due to the finite range of simulated production fluxes *N*_2_*/N*_*x*_ (see SI).

Even in the idealized case with perfect adaptation, the above class of control systems is predicted to achieve noise suppression in the ⟨*z*_1_⟩ ≫1, ⟨*z*_2_⟩≪ 1 regime, in which the linear noise approximation yields

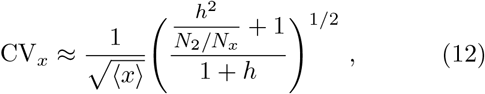

where 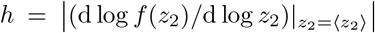 quantifies the non-linearity of the control input function. Significant noise suppression should thus be possible for the sensor-actuated control while conferring *perfect* adaptation as long as *h* < *N*_2_/*N*_*x*_. Indeed, exact numerical simulations verify that idealized sensor-actuated AIF systems can suppress noise significantly below open-loop levels (Fig. 4B). This corroborates that there is no fundamental trade-off between achieving robust adaptation of averages versus noise suppression with AIF control.

Prior work considered closed-loop antithetic integral ‘rein control’ systems in which *both Z*_1_ and *Z*_2_ affect the controlled system and established that the combined feedback can achieve lower noise than feedback through the reference species only [9]. However, the previously reported numerical data for such ‘rein control’ systems still exhibited a noise penalty compared to the corresponding open-loop fluctuations (see SI). Other work [56] demonstrated noise suppression for an alternative biomolecular integral control strategy proposed in [57]. However, as previously noted, this simpler strategy is unsuitable in the stochastic setting due the presence of an absorbing state for the controller species [57]. Moreover, if the ensemble average dynamics are analyzed in the stochastic regime without making a deterministic approximation as in [56], the simplified controller proposed in [57] does not confer perfect adaptation, even if a non-zero stationary state were attainable.

Minimizing Eq. 12 over all values of *h* predicts a bound on the noise suppression of the sensor-actuated AIF module

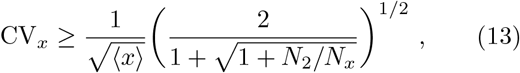

which is obeyed by our numerical explorations of idealized sensor-actuated AIF (SI).

Furthermore, the linear noise approximation predicts Eq. 13 constrains the sensor-actuated AIF system when the decay reactions given in Eq. 7 are included (see SI). Our sensitivity approximation predicts that the right side of Eq. 13 is achieved for 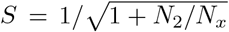 (see SI). Thus, for a fixed *N*_2_*/N*_*x*_, we predict both the minimum noise level and corresponding sensitivity are smaller for the controller with sensor-actuation than with reference-actuation. This suggests that the improved noise properties of sensor-actuated AIF vs. reference-actuated AIF extend to the non-idealized regime with dilution in their optimal noise regimes for a given sensor burden *N*_2_*/N*_*x*_.

Intriguingly, both Eq. 13 and Eq. 9 have the same asymptotic behaviour as the fundamental information theoretic bound that applies to any molecular feedback system in which a controlled species undergoes first order degradation and its abundance is read out through a sensing reaction with a linear rate [58]. This may indicate that noise suppression performance is bounded similarly for feedback systems regardless of their ability to achieve perfect adaptation for averages.

Note, the inability of reference-actuated AIF control to suppress noise in the idealized regime (Fig. 2B) is not due to the linear actuation rate of Eq. 4: When generalizing the idealized reference-actuated AIF module to allow for a non-linear control input function, the linear noise approximation still predicts increased noise compared to open-loop systems (see SI). Moreover, Eq. 13 predicts the sensor-actuated system can suppress fluctuations even in the linear regime *h* = 1. The drastically different noise properties of sensor-actuated and reference-actuated AIF modules thus seem due to their different control structure and not their different control rates.

## DISCUSSION

The term homeostasis was coined by Walter Cannon to describe how large environmental disturbances lead to only small internal changes in adaptive biological systems [59]. For example, steady state human core body temperature changes only around 1°C in response to changes in ambient temperature of 11°C, even in the absence of thermal insulation through clothing [60]. Our results show that models of *perfect* adaptation for homeostatic systems can be misleading because their behaviour can be singular: even in the infinitesimal limit, near-perfectly adaptive biological systems do not universally approach the behaviour of systems with perfect adaptation.

In particular, we challenge the notion that there exists a fundamental trade-off between achieving low sensitivity to external perturbations for average abundances and low variability around the average that has previously been proposed based on the analysis of perfectly adapting systems [1, 2, 32, 33]. In contrast to their idealized counterparts, we find that reference-actuated AIF systems can achieve significant noise suppression levels for even minute deviations from perfect adaptation, as long as long as cells can pay the corresponding energetic price. For example, we present a non-idealized realization of the AIF controller which reduces noise by over 80% while exhibiting a minuscule sensitivity of less than 0.02. The cost of combining such noise suppression performance with near perfect adaptation came in the form of producing ∼890,000 controller molecules per control molecule.

Our analysis of AIF control in which the sensor species, rather than the reference species, acts on the controlled species further corroborates the absence of a fundamental trade-off between adaptation and noise suppression We find that this atypical variant of the AIF module can suppress noise even in the idealized perfect adaptation regime without any secondary control structures. This may prove an important design consideration for synthetic biology applications in which increased complexity due to additional control circuits can be detrimental to performance: Similar, to how recent improvements of engineered genetic oscillators came through simplification rather than through additional control structures [61], the adaptation-conferring AIF control module devised by Briat, *et al*. [1] might be best left unadulterated – even when noise suppression performance is required.

Our work has several limitations. All presented results are based on the analysis of a reference open-loop system composed of a single species whose levels are Poisson distributed in the absence of control. Further, our analyses do not explicitly account for the cell-cycle and division. While our results establish that sensor-actuated AIF control has preferable noise properties compared to reference-actuated variants, future work is required to establish the underlying control theoretic reasons for the observed difference in noise. However, our results based on minimal AIF models provide insights into the limitations of robust biological control and make predictions for improving the performance of robustly adapting synthetic gene regulatory circuits that may be tested experimentally and theoretically for more complex systems.

## MATERIALS & METHODS

### Analytical Approximations

The linear noise approximation is a standard approximation technique for the analysis of non-linear stochastic reaction networks (e.g., see [34–37]). Here, we present a simple derivation of the approximation from stationary second-order moment invariants for discrete Markov processes [36, 62].

Consider general discrete stochastic processes where the state vector ***y*** = (*y*_0_, …, *y*_*N*−1_) is updated through transitions

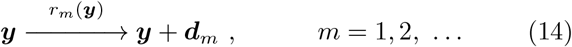

that change the level of a species *Y*_*i*_ by a discrete jump of size *d*_*im*_ with a state-dependent probability per unit time *r*_*m*_(***y***). To specify the stationary second-order moment invariants, following [62], we first specify several quantities to characterize the system. The total birth 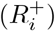 and death 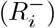 fluxes for a species *Y*_*i*_ are given by 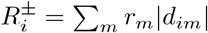and *d*_*im*_ < 0, respectively. Little’s law [62–64] from queuing theory then gives the average lifetime as

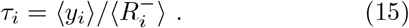

The average jump size ⟨*s*_*ij*_⟩, defined as the average change in the number of *Y*_*j*_ molecules as a *Y*_*i*_ molecule is made or degraded, is given by

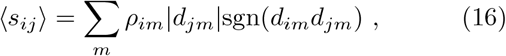

where *ρ*_*im*_ is the fraction of flux of species *Y*_*i*_ going through reaction *m*. At stationarity, any system of the form given by Eq. 14 satisfies

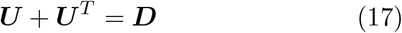

for the matrices with entries

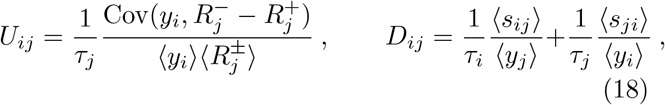

see [62]. In the limit of infinitesimal fluctuations, the birth-death fluxes 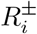 can be well-approximated by a first-order expansion

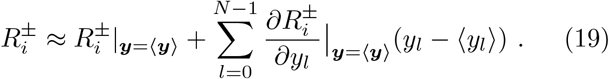

It follows that

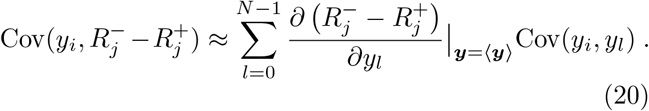

Straightforward manipulations then show that in the small noise limit, Eq. 17 is approximated by the con-tinuous Lyapunov equation

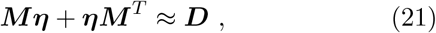

where ***η*** is the (approximate) stationary covariance matrix with entries *η*_*ij*_ = Cov(*y*_*i*_, *y*_*j*_)*/*(⟨*y*_*i*_⟩ ⟨*y*_*j*_⟩) and ***M*** is a ‘normalized Jacobian’ matrix with entries 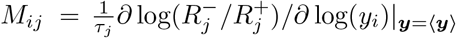 If −*M* is Hurwitz stable, then Eq. 21 permits a unique positive semi-definite solution [65]

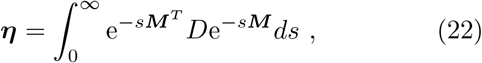

which we refer to as the linear noise approximation for the stationary covariance matrix.

### Numerical Simulations

Numerical simulations were run using the standard Gillespie algorithm [38, 39] implemented in C**++**11. Stationary joint probability distributions for the abundances of species were generated numerically by calculating the fraction of the total system time spent in each sampled state after each reaction occurred at least 10^6^ times. Under ergodic conditions, these long-time distributions converge to the unique stationary distribution representing both the temporal and ensemble statistics of the stochastic process. First-and second-order moments of the numerical distributions were calculated to generate the data used to validate our conjectured bounds. Analysis and plotting of numerical data was performed using scripts implemented in MATLAB R2020b. Simulations were included in the final analysis if they satisfied the first-and second-order moment invariants for stationary discrete Markov processes within 2% and 5% relative error, respectively (see SI). When evaluating the agreement of data with the conjectured bounds, the sampling error on edge cases was evaluated by simulating a given parameter set using at least three unique pseudo-random number generator seeds. See SI for a complete description of the error analysis and parameter sampling.

The sensitivity of the average *X*-levels to perturbations in the degradation rate *β*_*x*_ was determined numerically using a simple mid-point rule [66] to approximate the derivative of *(x)* with respect to *β*_*x*_:

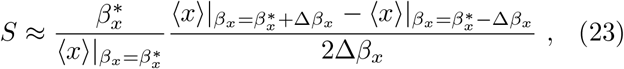

Where 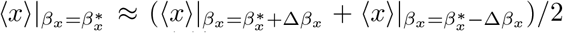. We approximate 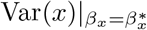 analogously to calculate the coefficient of variation corresponding to the numerical sensitivity coefficient. We generated our data using a relative perturbation size of 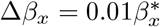 and worked in units of 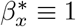.

### Data and code availability

Numerical simulation code and data will be available on Github along with the code for analysing this data and generating the figures for the manuscript.

## AUTHOR CONTRIBUTIONS

BK derived analytical results, generated numerical data, and analyzed the data. RR generated numerical data. BK and AH wrote the manuscript. All authors reviewed the manuscript and approved it for publication. The authors declare no competing interests.

## ACKNOWLEDGEMENTS

We thank Glenn Vinnicombe, David McMillen, and Zhe Tang for insightful comments that have helped us to significantly improve the manuscript. We thank Raymond Fan, Seshu Iyengar, and Euan Joly-Smith for many helpful discussions and suggestions throughout the work. We thank Johan Paulsson for bringing to our attention the etymology of the term homeostasis. This work was supported by the Natural Sciences and Engineering Research Council of Canada and a New Researcher Award from the University of Toronto Connaught Fund. BK gratefully acknowledges funding from the University of Toronto Faculty of Arts & Science Top Doctoral Fellowship.

