## SupplementaryInformation for "Noise properties of adaptation-conferring biochemical control modules"

#### This PDF file includes:

Supporting text
Figs. S1 to S5
Table S1
SI References

#### Other supporting materials for this manuscript include the following:

Wolfram Mathematica Notebook SupplementaryDerivations.nb

### Supporting Information Text

**S3 Supplementary figures and table** **15**

#### S1. Mathematical derivations

Here we present derivations of the main text equations and additional supporting derivations. We make use of `Wolfram Mathematica 12` throughout the derivations. The Wolfram Notebook Supplementary-Derivations.nb included with this Supplementary Text may be opened with the `Wolfram Mathematica` software itself or using the free `Wolfram Player` software (https://www.wolfram.com/player/).

**A. Linear Noise Lyapunov equation for the general three-component antithetic integral feedback topology.** Consider the system defined by the following stochastic transitions:

$$\begin{array}{cc} x \xrightarrow{f(x,z_1,z_2)} x+1 & x \xrightarrow{\beta_x x} x-1 \\ z_1 \xrightarrow{\mu} z_1+1 & z_1 \xrightarrow{\beta_1 z_1} z_1-1 \\ z_2 \xrightarrow{\theta x} z_2+1 & z_2 \xrightarrow{\beta_2 z_2} z_2-1 \\ \multicolumn{2}{c}{(z_1,\ z_2) \xrightarrow{\gamma z_1 z_2} (z_1-1,\ z_2-1)} \end{array} \tag{S1}$$

Each system considered in the subsequent derivations will be a special case of the above system.

Here we derive the Lyanunov equation for the linear noise approximation (see Materials and Methods) of the above generalized system's stationary covariance matrix. First we find the average lifetime of each species, which are given by Little's Law (1–3) (see Eq. 15) as follows:

$$\begin{aligned}\tau_x &= \frac{1}{\beta_x}\\ \tau_1 &= \frac{\langle z_1\rangle}{\beta_1\langle z_1\rangle + \gamma\langle z_1 z_2\rangle}\\ \tau_2 &= \frac{\langle z_2\rangle}{\beta_2\langle z_2\rangle + \gamma\langle z_1 z_2\rangle}\ .\end{aligned} \tag{S2}$$

Next, we define the elasticity parameters $h_x,\ h_1,\ h_2$ to quantify the nonlinearity of the actuation function $f(x, z_1, z_2)$ with respect to $x,\ z_1$ and $z_2$, respectively, as follows:

$$\begin{aligned}h_x &:= \frac{\partial f(x, z_1, z_2)}{\partial x}\Big|_{(x,z_1,z_2)=\langle(x,z_1,z_2)\rangle}\\ h_1 &:= \frac{\partial f(x, z_1, z_2)}{\partial z_1}\Big|_{(x,z_1,z_2)=\langle(x,z_1,z_2)\rangle}\\ h_2 &:= \frac{\partial f(x, z_1, z_2)}{\partial z_2}\Big|_{(x,z_1,z_2)=\langle(x,z_1,z_2)\rangle}\ .\end{aligned} \tag{S3}$$

Additionally, we define

$$\begin{aligned}E_1 &:= \frac{\gamma\langle z_1 z_2\rangle}{\beta_1\langle z_1\rangle + \gamma\langle z_1 z_2\rangle}\\ E_2 &:= \frac{\gamma\langle z_1 z_2\rangle}{\beta_2\langle z_2\rangle + \gamma\langle z_1 z_2\rangle}\end{aligned} \tag{S4}$$

which, respectively, specify the fraction of $Z_1$ molecules which eventually annihilate a $Z_2$ molecule's biological activity and the fraction of $Z_2$ molecules which eventually annihilate a $Z_1$ molecule's biological activity (3).

Let us work in a basis such that the state vector is $\boldsymbol{y} = (x,\ z_1,\ z_2)$. Then the normalized "Jacobian" $\boldsymbol{M}$ and diffusion matrix $\boldsymbol{D}$ needed to specify the linear noise approximation (4–7) for this system's stationary covariance matrix are given as follows:

$$\begin{aligned}\boldsymbol{M} &= \begin{pmatrix} \frac{1-h_x}{\tau_x} & -\frac{h_1}{\tau_x} & -\frac{h_2}{\tau_x}\\ 0 & \frac{1}{\tau_1} & \frac{E_1}{\tau_1}\\ -\frac{1}{\tau_2} & \frac{E_2}{\tau_2} & \frac{1}{\tau_2}\end{pmatrix}\\ \boldsymbol{D} &= \begin{pmatrix} \frac{2}{\tau_x\langle x\rangle} & 0 & 0\\ 0 & \frac{2}{\tau_1\langle z_1\rangle} & \frac{1}{2}\left(\frac{E_2}{\tau_2\langle z_1\rangle} + \frac{E_1}{\tau_1\langle z_2\rangle}\right)\\ 0 & \frac{1}{2}\left(\frac{E_2}{\tau_2\langle z_1\rangle} + \frac{E_1}{\tau_1\langle z_2\rangle}\right) & \frac{2}{\tau_2\langle z_2\rangle}\end{pmatrix}\ .\end{aligned} \tag{S5}$$

The approximate stationary covariance matrix $\boldsymbol{\eta}$ for this system is then obtained by solving the following Lyapunov equation for the above matrices:

$$\boldsymbol{M}\boldsymbol{\eta} + \boldsymbol{\eta}\boldsymbol{M}^T \approx \boldsymbol{D}\ . \tag{S6}$$

See Materials and Methods for a derivation of the linear noise approximation specified above.

We will also use the following stationary flux balance relations throughout the subsequent sections:

$$\begin{aligned}\langle f(x,z_1,z_2)\rangle &= \beta_x\langle x\rangle\\ \mu &= \gamma\langle z_1 z_2\rangle + \beta_1\langle z_1\rangle\\ \theta\langle x\rangle &= \gamma\langle z_1 z_2\rangle + \beta_2\langle z_2\rangle\ ,\end{aligned} \tag{S7}$$

These are obtained from the ensemble average dynamics for this system, which follow directly from the Master Equation (4). For the system given by Eq. S1, these are given by

$$\begin{aligned}\frac{\mathrm{d}\mathbb{E}[x](t)}{\mathrm{d}t} &= \mathbb{E}[f(z_2)](t) - \beta_x\mathbb{E}[x](t)\\ \frac{\mathrm{d}\mathbb{E}[z_1](t)}{\mathrm{d}t} &= \mu - \gamma\mathbb{E}[z_1z_2](t) - \beta_1\mathbb{E}[z_1](t)\\ \frac{\mathrm{d}\mathbb{E}[z_2](t)}{\mathrm{d}t} &= \theta\mathbb{E}[x](t) - \gamma\mathbb{E}[z_1z_2](t) - \beta_2\mathbb{E}[z_2](t)\ .\end{aligned} \tag{S8}$$

At stationarity the time derivatives of the ensemble averages vanish resulting in Eq. S7, where we have replaced ensemble averages with time averages, as we consider ergodic systems.

**B. Derivation of Eq. 6.** Here we consider the minimal idealized reference-actuated antithetic integral feedback (AIF) system with a linear actuation rate. The transitions for this system are given by Eq. S1 with $\beta_1=\beta_2=0$ and $f(x,z_1,z_2)=kz_1$. We approximate the stationary covariance matrix $\boldsymbol{\eta}$ for this system using the linear noise approximation (see Materials and Methods) for the matrices specified by Eq. S5 with $E_1=E_2=1$, $h_x=h_2=0$, and $h_1=1$. We solve the resulting Lyapunov equation using the `Solve[]` function in `Wolfram Mathematica 12`. This leads to an approximation for the coefficient of variation for the controlled species through the relation $\mathrm{CV}_x=\sqrt{\eta_{xx}}$, which is given as follows:

$$\mathrm{CV}_x^2 \approx \frac{1}{\psi}\Bigg(\frac{2\tau_2\tau_1^2+2\tau_2^2\tau_1+2\tau_1^2\tau_x+4\tau_2\tau_1\tau_x+2\tau_2^2\tau_x}{\langle x\rangle}+\frac{2\tau_2^2\tau_1+\tau_1^2\tau_2+\tau_1^2\tau_x+\tau_2\tau_1\tau_x}{\langle z_1\rangle}+\frac{\tau_1\tau_2^2+\tau_2^2\tau_x+\tau_1\tau_2\tau_x}{\langle z_2\rangle}\Bigg)\ , \tag{S9}$$

where

$$\psi = 2\left(\tau_2\tau_1^2+\tau_2^2\tau_1+\tau_1^2\tau_x+\tau_2\tau_1\tau_x+\tau_2^2\tau_x\right)\ . \tag{S10}$$

The stationary flux balance relations (see Eq. S7) for $Z_1$ and $Z_2$ ensure $\mu=\gamma\langle z_1z_2\rangle=\theta\langle x\rangle$ such that Eq. S2 implies $\tau_1=\langle z_1\rangle/(\theta\langle x\rangle)$ and $\tau_2=\langle z_2\rangle/(\theta\langle x\rangle)$ for this system. Substituting the above identity for $\tau_2$ and taking the limit of $\langle z_2\rangle\to\infty$ of the approximate $\mathrm{CV}_x$ leads to (see Wolfram Notebook SupplementaryDerivations.nb)

$$\lim_{\langle z_2\rangle\to\infty}\mathrm{CV}_x = \left(\frac{1}{\langle x\rangle}+\frac{\tau_1}{\langle z_1\rangle(\tau_1+\tau_x)}\right)^{1/2}\ . \tag{S11}$$

Furthermore, we verify using the `Reduce[]` function in `Wolfram Mathematica 12` that this limit bounds all other physical solutions for the approximate $\mathrm{CV}_x$. The linear noise approximation thus predicts

$$\mathrm{CV}_x \geq \left(\frac{1}{\langle x\rangle}+\frac{1}{\langle z_1\rangle(1+\tau_x/\tau_1)}\right)^{1/2}\ . \tag{S12}$$

The average number of birth events of $Z_2$ molecules during the average lifetime $\tau_x$ of the controlled species $X$ is given by $N_2=\theta\langle x\rangle\tau_x$ and the average number of birth events of $X$ molecules during the average lifetime $\tau_x$ of the controlled species $X$ is given by $N_x = k\langle z_1\rangle\tau_x$, where $\tau_x$ is given by Eq. S2. The stationary flux balance relations (see Eq. S7) for $X$ ensures $k\langle z_1\rangle = \langle x\rangle/\tau_x$ such that $N_x = \langle x\rangle$. Thus,

$$\langle z_1\rangle\tau_x/\tau_1 = \langle x\rangle N_2/N_x \ . \qquad \text{[S13]}$$

Combining this with Eq. S12 and pulling a factor of $1/\sqrt{\langle x\rangle}$ outside the square root results in the inequality denoted as Eq. 6 in the main text.

Using the stationary flux balance relation for $X$, Eq. S13 can equivalently be written

$$\tau_1 = \frac{1}{N_2/N_x}\frac{1}{k} \qquad \text{[S14]}$$

Thus, for a fixed $N_2/N_x$, $\tau_1$ can only be increased by decreasing $k$.

#### C. Sensitivity of reference-actuated antithetic integral feedback systems with dilution/degradation.

Here we consider the minimal reference-actuated AIF system including control species dilution/degradation with a linear actuation rate. The transitions for this system are given by Eq. S1 with $f(x, z_1, z_2) = kz_1$. For this system the general flux balance relations of Eq. S7 reduce to

$$\begin{aligned} k\langle z_1\rangle &= \beta_x\langle x\rangle \\ \mu &= \gamma\langle z_1z_2\rangle + \beta_1\langle z_1\rangle \\ \theta\langle x\rangle &= \gamma\langle z_1z_2\rangle + \beta_2\langle z_2\rangle \ , \end{aligned} \qquad \text{[S15]}$$

This system of equations is not algebraically closed due to the presence of the second-order moment $\langle z_1z_2\rangle$. However, by definition of the normalized covariance $\eta_{12}$ (see Materials and Methods), we have the following identity:

$$\langle z_1z_2\rangle = (\eta_{12} + 1)\langle z_1\rangle\langle z_2\rangle \ . \qquad \text{[S16]}$$

We use this identity to approximate $\langle z_1z_2\rangle$ by substituting the linear noise approximation for $\eta_{12}$. This is obtained by solving the linear noise approximation (see Eq. 21) for the matrices specified by Eq. S5 with $h_x = h_2 = 0$, and $h_1 = 1$. This leads to the following approximation:

$$\begin{aligned} \eta_{12} \approx \frac{1}{\psi}\Bigg(&\frac{1}{\langle z_1\rangle}(E_1E_2\tau_1\tau_x^2 + 2E_1\tau_2\tau_1\tau_x + E_2\tau_2\tau_1^2 + E_2\tau_1^2\tau_x + E_2\tau_1\tau_x^2 + E_2\tau_2\tau_1\tau_x - E_1E_2^2\tau_1\tau_x^2 - \\ &2\tau_2\tau_1^2 - 2\tau_1^2\tau_x - 2E_1E_2\tau_2\tau_1\tau_x) + \frac{2E_1\tau_1\tau_x^2 + 2E_1\tau_2\tau_x^2 + 2E_1\tau_1\tau_2\tau_x}{\langle x\rangle} + \\ &\frac{E_1^2\tau_2\tau_x^2 + E_1\tau_1\tau_2^2 + E_1\tau_2^2\tau_x + E_1\tau_2\tau_x^2 + E_1\tau_1\tau_2\tau_x - E_1^2E_2\tau_2\tau_x^2}{\langle z_2\rangle}\Bigg) \ , \end{aligned} \qquad \text{[S17]}$$

where

$$\begin{aligned} \psi = 2(1 + E_1 - E_1E_2)(&E_1E_2\tau_1\tau_x^2 + E_1E_2\tau_2\tau_x^2 + E_1\tau_2\tau_1\tau_x - \\ &\tau_2\tau_1^2 - \tau_2^2\tau_1 - \tau_1^2\tau_x - \tau_1\tau_x^2 - 2\tau_2\tau_1\tau_x - \tau_2\tau_x^2 - \tau_2^2\tau_x) \ . \end{aligned} \qquad \text{[S18]}$$

It straightforward to show by combining Eq. S2, Eq. S4, and the flux balance relations for $Z_1$ and $Z_2$ in Eq. S15 that the following identities hold:

$$\begin{aligned}\langle x\rangle &= \frac{\mu}{\theta}\frac{E_1}{E_2}\\ \langle z_1\rangle &= \frac{\mu}{\beta_1}(1-E_1)\\ \langle z_2\rangle &= \frac{\mu}{\beta_2}\frac{E_1}{E_2}(1-E_2)\\ \langle z_1 z_2\rangle &= \frac{\mu}{\gamma}E_1\\ \tau_1 &= \frac{1-E_1}{\beta_1}\\ \tau_2 &= \frac{1-E_2}{\beta_2}\\ \tau_x &= \frac{1}{\beta_x}\ .\end{aligned} \qquad [S19]$$

We then combine Eqs. S16, S17 and substitute the above identities into the resulting expression and the flux balance relation for $X$ in Eq. S15. Differentiating both sides of the resulting expressions with respect to $\beta_x$ results in a system of two equations in the unknowns $\partial E_1/\partial\beta_x$ and $\partial E_2/\partial\beta_x$, which we solve using the `Solve[]` function in `Wolfram Mathematica 12` (see Wolfram Notebook SupplementaryDerivations.nb). The approximate expressions for $\partial E_1/\partial\beta_x$ and $\partial E_2/\partial\beta_x$ are then used to obtain an approximation for $\partial\langle x\rangle/\partial\beta_x$ through the identity

$$\frac{\partial\langle x\rangle}{\beta_x} = \frac{\mu}{\theta}\frac{1}{E_2^2}\left(E_2\frac{\partial E_1}{\partial\beta_x} - E_1\frac{\partial E_2}{\partial\beta_x}\right)\ , \qquad [S20]$$

which follows by differentiating the expression for $\langle x\rangle$ in Eq. S19. The sensitivity coefficient defined in Eq. 8 can equivalently be written

$$S = \frac{\beta_x}{\langle x\rangle}\frac{\partial\langle x\rangle}{\partial\beta_x}\ . \qquad [S21]$$

The sensitivity coefficient is thus approximated by substituting the approximation for $\partial\langle x\rangle/\partial\beta_x$ and the $\langle x\rangle$ identity in Eq. S19. The resulting expression is algebraically extremely involved but still tractable using `Wolfram Mathematica`. We have thus included the final expression in the attached Wolfram Notebook SupplementaryDerivations.nb.

**D. Derivation of Eqs. 9 and 10.** Here we consider the minimal reference-actuated AIF system including control species dilution/degradation with a linear actuation rate. The transitions for this system are given by Eq. S1 with $f(x, z_1, z_2) = kz_1$. We approximate the stationary covariance matrix $\boldsymbol{\eta}$ for this system using the linear noise approximation (see Materials and Methods) for the matrices specified by Eq. S5 with $h_x = h_2 = 0$, and $h_1 = 1$. We solve the resulting Lyapunov equation using the `Solve[]` function in `Wolfram Mathematica 12`. This leads to an approximation for the coefficient of variation for the controlled species through the relation $\mathrm{CV}_x = \sqrt{\eta_{xx}}$ (see Wolfram Notebook SupplementaryDerivations.nb). Let us define the Fano factor as a measure of the noise the controlled system exhibits relative to the open-loop Poisson noise

$$F_x := \frac{\mathrm{CV}_x^2}{1/\langle x\rangle}\ , \qquad [S22]$$

where $\mathrm{CV}_x$ is approximated as described above. We derive a lower bound on $F_x$, which corresponds to a lower bound on $\mathrm{CV}_x$ relative to Poisson fluctuations, as desired. This is simply to avoid introducing square roots, which simplifies the calculations. First, we show the following inequality holds in the $F_x < 1$ regime:

$$\begin{aligned} F_x &\geq \lim_{\beta_1/\beta_x,\ \beta_1/\theta \to \infty} F_x\Big|_{\beta_2 = c\beta_1} \\ &= \underbrace{\frac{E_1^2 E_2^3 \theta\tau_x - 2E_1E_2^2\theta\tau_x + E_1 + E_2\theta\tau_x}{E_2\theta\tau_x\left(E_1^2(E_2-1)E_2 - 2E_1E_2 + E_1 + 1\right)}}_{=:B(E_1,E_2,\theta\tau_x)}\ , \end{aligned} \qquad \text{[S23]}$$

where $c := \beta_2/\beta_1$. This follows by first showing that the only critical value of $\beta_1$ satisfying $\partial F_x|_{\beta_2=c\beta_1}/\partial\beta_1 = 0$ corresponds to a maximum of $F_x|_{\beta_2=c\beta_1}$ for given values of $E_1$, $E_2$, $\theta\tau_x$ and $c$ (see Wolfram Notebook SupplementaryDerivations.nb). We then show $\lim_{\beta_1/\beta_x,\ \beta_1/\theta\to 0} F_x|_{\beta_2=c\beta_1} = 1$ (see Wolfram Notebook SupplementaryDerivations.nb). $F_x|_{\beta_2=c\beta_1}$ can be expressed as a ratio of quadratics in $\beta_1$ for which the roots of the denominator are strictly negative for allowed values of $E_1$, $E_2$, $\tau_x$ and $c$ (see Wolfram Notebook SupplementaryDerivations.nb). Thus, $F_x|_{\beta_2=c\beta_1}$ is continuous with respect to $\beta_1 > 0$. It follows that $\lim_{\beta_1/\beta_x,\ \beta_1/\theta\to\infty} F_x\Big|_{\beta_2=c\beta_1}$ minimizes $F_x$ in the $F_x < 1$ regime. For given values of $E_2$ and $\theta\tau_x$, the following is the unique critical $E_1 \leq 1$ value satisfying $\partial \lim_{\beta_1/\beta_x,\ \beta_1/\theta\to\infty} F_x\Big|_{\beta_2=c\beta_1}/\partial E_1 = 0$:

$$E_1^*(E_2, \theta\tau_x) = \frac{E_2^2\theta\tau_x - \sqrt{E_2(E_2\theta\tau_x + E_2 - 1)}}{E_2\left(E_2^2\theta\tau_x + E_2 - 1\right)}\ , \qquad \text{[S24]}$$

which corresponds to a minimizer of $B(E_1, E_2, \theta\tau_x)$ for given values of $E_2$ and $\theta\tau_x$ (see Wolfram Notebook SupplementaryDerivations.nb). Combined with Eq. S23, we thus have

$$F_x \geq B(E_1^*(E_2,\theta\tau_x), E_2, \theta\tau_x)\ . \qquad \text{[S25]}$$

Furthermore, setting $E_2 = 1$, we find

$$B(E_1^*(1,\theta\tau_x), 1, \theta\tau_x) = \frac{2\sqrt{\theta\tau_x} - 1}{\theta\tau_x}\ . \qquad \text{[S26]}$$

We verify using the `Reduce[]` function in `Wolfram Mathematica 12` that in the $F_x < 1$ regime the following inequality holds:

$$B(E_1^*(E_2,\theta\tau_x), E_2, \theta\tau_x) \geq \frac{2\sqrt{\theta\tau_x} - 1}{\theta\tau_x}\ . \qquad \text{[S27]}$$

Combined with Eq. S25, we have

$$F_x \geq \frac{2\sqrt{\theta\tau_x} - 1}{\theta\tau_x}\ . \qquad \text{[S28]}$$

Eq. 9 is then obtained by identifying $N_2/N_x = \theta\tau_x$ and rewriting the resulting inequality in terms of $\mathrm{CV}_x$ via the definition Eq. S22.

We now derive Eq. 10. First recall the definition of $\mathrm{CV}_x^{\min}$ from main text Eq. 9:

$$\mathrm{CV}_x^{\min} := \left(\frac{2\sqrt{N_2/N_x} - 1}{N_2/N_x}\right)^{1/2}\ , \qquad \text{[S29]}$$

which constrains $\mathrm{CV}_x$ according to $\mathrm{CV}_x \geq \mathrm{CV}_x^{\min}$, where $\mathrm{CV}_x \equiv \sqrt{F_x}$ is obtained via the linear noise approximation. As revealed by the derivation of Eq. S28 above, $\mathrm{CV}_x$ approaches $\mathrm{CV}_x^{\min}$ in the limit of $\beta_1/\beta_x$, $\beta_1/\theta \to \infty$ with $\beta_2 \propto \beta_1$ for $E_1 = E_1^*(E_2, \theta\tau_x)$ and $E_2 = 1$, where $E_1^*(E_2,\theta\tau_x)$ is defined by

Eq. S24. Taking the $E_2 \to 1$ limit of the approximate sensitivity coefficient $S$ derived in Sect. S1.C gives (see Wolfram Notebook SupplementaryDerivations.nb)

$$\lim_{E_2 \to 1} S = 1 - E_1 \ . \qquad [S30]$$

The sensitivity value $S = S^*$ which minimizes the approximate $\mathrm{CV}_x$ for a given $\theta\tau_x$ is then calculated by substituting $E_1 = E_1^*(1, \theta\tau_x)$, where $E_1^*(E_2, \theta\tau_x)$ is given by Eq. S24, resulting in

$$S^* = 1/\sqrt{N_2/N_x} \ , \qquad [S31]$$

where we have invoked the identity $N_2/N_x = \theta\tau_x$. Eq. 10 then follows from substituting Eq. S31 into Eq. S29.

**E. Derivation of Eqs. 12 and 13.** Here we consider the minimal idealized sensor-actuated AIF system. The transitions for this system are given by Eq. S1 with $\beta_1 = \beta_2 = 0$ and $f(x, z_1, z_2) = f(z_2)$, for some $f : \mathbb{Z} \to \mathbb{R}_{\geq 0}$ characterized by $h_2 = (\mathrm{d}\log f(z_2)/\mathrm{d}\log z_2)|_{z_2=\langle z_2\rangle} < 0$. We approximate the stationary covariance matrix $\boldsymbol{\eta}$ for this system using the linear noise approximation (see Materials and Methods) for the matrices specified by Eq. S5 with $E_1 = E_2 = 1$, $h_x = h_1 = 0$. We solve the resulting Lyapunov equation using the `Solve[]` function in `Wolfram Mathematica 12`. This leads to an approximation for the coefficient of variation for the controlled species through the relation $\mathrm{CV}_x = \sqrt{\eta_{xx}}$, which is given as follows:

$$\mathrm{CV}_x^2 \approx \frac{1}{\psi}\left(\frac{2\tau_2\tau_1^2 + 2\tau_2^2\tau_1 + 2\tau_1^2\tau_x + 4\tau_2\tau_1\tau_x + 2\tau_2^2\tau_x - 2h_2\tau_2\tau_1^2}{2\langle x\rangle} - \frac{h_2\tau_2\tau_1^2 + h_2\tau_1^2\tau_x + h_2\tau_2\tau_1\tau_x}{2\langle z_1\rangle} + \frac{2h_2^2\tau_2\tau_1^2 - h_2\tau_2^2\tau_1 - h_2\tau_2\tau_1\tau_x - h_2\tau_2^2\tau_x}{2\langle z_2\rangle}\right) , \qquad [S32]$$

where

$$\psi = 2(\tau_2\tau_1^2 + \tau_2^2\tau_1 + \tau_1^2\tau_x + 2\tau_2\tau_1\tau_x + \tau_2^2\tau_x - h_2\tau_1^2\tau_x - h_2\tau_2\tau_1^2) \ . \qquad [S33]$$

The stationary flux balance relations (see Eq. S7) for $Z_1$ and $Z_2$ ensures $\mu = \gamma\langle z_1 z_2\rangle = \theta\langle x\rangle$ such that Eq. S2 implies $\tau_1 = \langle z_1\rangle/(\theta\langle x\rangle)$ and $\tau_2 = \langle z_2\rangle/(\theta\langle x\rangle)$ for this system. Substituting the above lifetime identities and taking the limit of $\langle z_1\rangle \to \infty$ and $\langle z_2\rangle \to 0$ of the approximate $\mathrm{CV}_x$ leads to (see Wolfram Notebook SupplementaryDerivations.nb)

$$\lim_{\langle z_1\rangle \to \infty} \lim_{\langle z_2\rangle \to 0} \mathrm{CV}_x = \frac{1}{\sqrt{\langle x\rangle}}\left(\frac{\frac{h_2^2}{\theta\tau_x} + 1}{1 - h_2}\right)^{1/2} . \qquad [S34]$$

Eq. 12 is obtained by identifying $N_2/N_x = \theta\tau_x$ and defining $h := |h_2|$.

Furthermore, we verify using the `Reduce[]` function in `Wolfram Mathematica 12` that this limit bounds all other physical solutions for the approximate $\mathrm{CV}_x$ in the $\mathrm{CV}_x < 1$ regime. Thus, we have the prediction

$$\mathrm{CV}_x \geq \frac{1}{\sqrt{\langle x\rangle}} \min\left\{\left(\frac{\frac{h_2^2}{N_2/N_x} + 1}{1 - h_2}\right)^{1/2}, \ 1\right\} . \qquad [S35]$$

Minimizing the right side of this inequality over all $h_2$ yields two roots for the critical $h_2$ value. The negative root is given

$$h_2^* = 1 - \sqrt{1 + N_2/N_x} \ . \qquad [S36]$$

and corresponds to a minimizer of the bound in Eq. S35. Substituting Eq. S36 in Eq. S35 and simplifying then gives Eq. 13.

**F. Linear noise analysis of a minimal antithetic integral rein control system.** Here we consider the minimal idealized sensor-actuated AIF system with an additional control input that depends on the reference species. This control strategy with two simultaneous control inputs from the AIF module has previously been termed antithetic integral rein control (8). The transitions for this system are given by Eq. S1 with $\beta_1 = \beta_2 = 0$ and $f(x, z_1, z_2) = f(z_1, z_2) = f_1(z_1) + f_2(z_2)$, for some $f_1, f_2 : \mathbb{Z} \to \mathbb{R}_{\geq 0}$ characterized by $h_1 = (\partial \log f(z_1, z_2)/\partial \log z_1)|_{(z_1,z_2)=\langle(z_1,z_2)\rangle} > 0$, $h_2 = (\partial \log f(z_1, z_2)/\partial \log z_2)|_{(z_1,z_2)=\langle(z_1,z_2)\rangle} < 0$. Note, the sensor species influences the controlled species via it's degradation rate in the antithetic integral rein controller defined in (8). However, the linear noise approximation for such a system is equivalent to one where the sensor represses controlled species levels by decreasing the birth rate of controlled species molecules, as specified here. We approximate the stationary covariance matrix $\boldsymbol{\eta}$ for this system using the linear noise approximation (see Materials and Methods) for the matrices specified by Eq. S5 with $E_1 = E_2 = 1$, $h_x = 0$. We solve the resulting Lyapunov equation using the `Solve[]` function in `Wolfram Mathematica 12`. This leads to an approximation for the coefficient of variation for the controlled species through the relation $\mathrm{CV}_x = \sqrt{\eta_{xx}}$, which is given as follows:

$$\mathrm{CV}_x^2 \approx \frac{1}{\psi}\Bigg(\frac{2\tau_2\tau_1^2 + 2\tau_2^2\tau_1 + 2\tau_1^2\tau_x + 4\tau_2\tau_1\tau_x + 2\tau_2^2\tau_x - 2h_2\tau_2\tau_1^2}{\langle x\rangle} + \frac{h_1\tau_2\tau_1^2 + h_1h_2\tau_2\tau_1^2 + 2h_1^2\tau_2^2\tau_1 + h_1\tau_1^2\tau_x + h_1\tau_2\tau_1\tau_x - h_2\tau_2\tau_1^2 - h_2\tau_1^2\tau_x - h_2\tau_2\tau_1\tau_x}{\langle z_1\rangle} + \frac{2h_2^2\tau_2\tau_1^2 + h_1\tau_2^2\tau_1 + h_1h_2\tau_2^2\tau_1 + h_1\tau_2\tau_1\tau_x + h_1\tau_2^2\tau_x - h_2\tau_2^2\tau_1 - h_2\tau_2^2\tau_x - h_2\tau_2\tau_1\tau_x}{\langle z_2\rangle}\Bigg)\,, \quad \text{[S37]}$$

where

$$\psi = 2(\tau_2\tau_1^2 + \tau_2^2\tau_1 + \tau_1^2\tau_x + 2\tau_2\tau_1\tau_x + \tau_2^2\tau_x - h_2\tau_2\tau_1^2 - h_2\tau_1^2\tau_x - h_1\tau_2\tau_1\tau_x)\,. \quad \text{[S38]}$$

S37 obeys the same limit as Eq. S34, which suggests noise suppression is attainable by the antithetic integral rein controller. However, in the $\mathrm{CV}_x < 1$ regime, the linear noise approximation additionally predicts that $\mathrm{CV}_x$ for the idealized antithetic integral rein control system is bounded from below by $\mathrm{CV}_x$ for the idealized sensor-actuated AIF system for equal averages, $h_2$, and $N_2/N_x$ (see Wolfram Notebook SupplementaryDerivations.nb). Furthermore, the results of (8) establish the antithetic integral rein controller can achieve much smaller settling-times than than a corresponding reference-actuated system. The transient dynamics of sensor-actuated systems (without an additional reference input) have not been investigated. Future work should investigate the transient dynamics of systems with only sensor-actuation.

While this analysis predicts that the antithetic integral rein control system can suppress noise relative to the system without the rein control module, existing studies only compared the fluctuations of an antithetic integral rein control system to the fluctuations of a corresponding reference-actuated system (8). Here we evaluate whether the numerical results of (8) confirm this prediction. The open-loop system considered in (8) is a two-step constitutive gene expression model with the following stochastic transitions:

$$r \xrightarrow{k_r} r+1 \qquad r \xrightarrow{\gamma_r r} r-1$$
$$p \xrightarrow{k_p p} p+1 \qquad p \xrightarrow{\gamma_p p} p-1\,, \quad \text{[S39]}$$

where $r$, $p$ denote the abundance of an mRNA species $R$ and its corresponding protein species $P$. Our one-component open-loop system can be regarded as the $\gamma_r \gg \gamma_p$ limit of this system with $X \equiv P$. Since this is a linear system, the linear noise approximation (see Materials and Methods) for the stationary covariance matrix $\boldsymbol{\eta}$ is exact. As previously derived (6), the normalized variance for the protein abundance is given

$$\eta_{pp} = \frac{1}{\langle p \rangle} + \frac{\tau_r}{\tau_p + \tau_r}\frac{1}{\langle r \rangle} \ , \qquad \text{[S40]}$$

where $\tau_r = 1/\gamma_r$, $\tau_p = 1/\gamma_p$. The stationary flux balance relation for the protein species ensures $k_p\langle r \rangle = \gamma_p\langle p \rangle$. Setting the open-loop protein average to $\langle p \rangle = \mu/\theta$ to match the closed-loop system, we then have $\langle r \rangle = \gamma_p\mu/(k_p\theta)$. Note, this corresponds to $k_r = \gamma_r\gamma_p\mu/(k_p\theta)$. Thus, the open-loop variance is given as

$$\mathrm{Var}(p) \equiv \eta_{pp}\langle p \rangle^2 = \frac{\mu}{\theta}\left(1 + \frac{k_p}{\gamma_p + \gamma_r}\right) \ . \qquad \text{[S41]}$$

Numerical stationary closed-loop variance values exceeding 50 for the open-loop system S39 under antithetic integral rein control for two different parameter sets are reported in (8). In both cases $\mu = 100$ and $\theta = \gamma_r = k_p = 5$. The first parameter set additionally corresponds to $\gamma_p = 5$, which leads to an open-loop variance of 30. The second parameter set additionally corresponds to $\gamma_p = 0.1$, which leads to an open-loop variance of $\approx 40$. Thus, the stationary closed-loop fluctuation values reported in (8) were larger than open-loop levels. However, the linear noise approximation derived for this system above predicts that noise suppression should be possible in some regimes of the 'rein controller', as intuitively expected for a control system that uses two control inputs, one of which we already showed to be able to suppress noise.

**G. Derivation of approximate sensitivity coefficient and minimum noise prediction for the sensor-actuated antithetic integral feedback system with dilution/degradation.** Here we consider the minimal reference-actuated AIF system including control species dilution/degradation. The transitions for this system are given by Eq. S1 with $f(x, z_1, z_2) = f(z_2)$, for some $f : \mathbb{Z} \to \mathbb{R}_{\geq 0}$ characterized by $h_2 = (\mathrm{d}\log f(z_2)/\mathrm{d}\log z_2)|_{z_2=\langle z_2 \rangle} < 0$. For this system the general flux balance relations of Eq. S7 reduce to

$$\begin{aligned} \langle f(z_2) \rangle &= \beta_x\langle x \rangle \\ \mu &= \gamma\langle z_1 z_2 \rangle - \beta_1\langle z_1 \rangle \\ \theta\langle x \rangle &= \gamma\langle z_1 z_2 \rangle - \beta_2\langle z_2 \rangle \ , \end{aligned} \qquad \text{[S42]}$$

Under the linear noise approximation we take $\langle f(z_2) \rangle \approx f(\langle z_2 \rangle)$. Upon invoking this, an identical procedure as in Sect. S1.C can be followed to derive an approximation for the sensitivity coefficient for this system. See Wolfram Notebook SupplementaryDerivations.nb for further details and the final expression. Moreover, an identical procedure to that in Sect. S1.D reveals that the analogous equations to S29 and S31 for this system are

$$\mathrm{CV}_x^{\min} = \frac{1}{\sqrt{\langle x \rangle}}\left(\frac{2}{1 + \sqrt{1 + N_2/N_x}}\right)^{1/2} \qquad \text{[S43]}$$

and

$$S^* = 1/\sqrt{1 + N_2/N_x} \ , \qquad \text{[S44]}$$

respectively. See Wolfram Notebook SupplementaryDerivations.nb for the full derivation.

**H. Linear noise analysis of idealized reference-actuated antithetic integral feedback system with nonlinear actuation.** Here we consider the minimal idealized reference-actuated antithetic integral feedback (AIF) system with a nonlinear actuation rate. The transitions for this system are given by Eq. S1 with $\beta_1 = \beta_2 = 0$ and $f(x, z_1, z_2) = f(z_1)$, for some $f : \mathbb{Z} \to \mathbb{R}_{\geq 0}$ characterized by $h_1 = (\mathrm{d}\log f(z_1)/\mathrm{d}\log z_1)|_{z_1=\langle z_1 \rangle} > 0$. We approximate the stationary covariance matrix $\boldsymbol{\eta}$ for this system using the linear noise approximation (see Materials and Methods) for the matrices specified by Eq. S5 with $E_1 = E_2 = 1$, $h_x = h_2 = 0$. We solve the resulting Lyapunov equation using the `Solve[]`

function in `Wolfram Mathematica 12`. This leads to an approximation for the coefficient of variation for the controlled species through the relation $\mathrm{CV}_x = \sqrt{\eta_{xx}}$, which is given as follows:

$$\mathrm{CV}_x^2 \approx \frac{1}{\psi}\Bigg(\frac{2\left(\tau_1+\tau_2\right)\left(\tau_2\tau_x+\tau_1\left(\tau_2+\tau_x\right)\right)}{\langle x\rangle}+\frac{h_1\tau_2\left(\tau_2\tau_x+\tau_1\left(\tau_2+\tau_x\right)\right)}{\langle z_2\rangle}+ \frac{h_1\tau_1\left(\tau_2\left(2h_1\tau_2+\tau_x\right)+\tau_1\left(\tau_2+\tau_x\right)\right)}{\langle z_1\rangle}\Bigg)\,, \quad [\mathrm{S45}]$$

where

$$\psi = 2\left(\tau_2\tau_1^2+\tau_2^2\tau_1+\tau_1^2\tau_x+2\tau_2\tau_1\tau_x+\tau_2^2\tau_x-h_1\tau_2\tau_1\tau_x\right)\,. \quad [\mathrm{S46}]$$

We verify using the `Reduce[]` function in `Wolfram Mathematica 12` that Eq. S45 predicts a noise penalty (see Wolfram Notebook SupplementaryDerivations.nb), i.e.,

$$\mathrm{CV}_x \geq \frac{1}{\sqrt{\langle x\rangle}}\,. \quad [\mathrm{S47}]$$

The linear noise approximation thus predicts that the noise penalty of the reference-actuated AIF applies to both linear as well as nonlinear actuation functions.

**I. Linear noise analysis of idealized reference-actuated antithetic integral feedback system with additional negative feedback input.** Here we consider the minimal idealized reference-actuated AIF system with a linear actuation rate and an additional negative self-feedback on the controlled species $X$. The transitions for this system are given by Eq. S1 with $\beta_1=\beta_2=0$ and $f(x,z_1,z_2)=f(x,z_1)=kz_1+g(x)$, for some $g:\mathbb{Z}\to\mathbb{R}_{\geq 0}$ characterized by $h_x = (\partial\log f(x,z_1)/\partial\log x)|_{(x,z_1)=\langle(x,z_1)\rangle}<0$. We approximate the stationary covariance matrix $\boldsymbol{\eta}$ for this system using the linear noise approximation (see Materials and Methods) for the matrices specified by Eq. S5 with $E_1=E_2=1$, $h_2=0$, and $h_1=1$. We solve the resulting Lyapunov equation using the `Solve[]` function in `Wolfram Mathematica 12`. This leads to an approximation for the coefficient of variation for the controlled species through the relation $\mathrm{CV}_x=\sqrt{\eta_{xx}}$, which is given as follows:

$$\mathrm{CV}_x^2 \approx \frac{1}{\psi}\Bigg(\frac{2\tau_2\tau_1^2+2\tau_2^2\tau_1+2\tau_1^2\tau_x+4\tau_2\tau_1\tau_x+2\tau_2^2\tau_x-2\tau_2\tau_1^2h_x-2\tau_2^2\tau_1h_x}{\langle x\rangle}+ \frac{2\tau_2^2\tau_1+\tau_1^2\tau_2+\tau_1^2\tau_x+\tau_2\tau_1\tau_x-\tau_2\tau_1^2h_x}{\langle z_1\rangle}+\frac{\tau_1\tau_2^2+\tau_2^2\tau_x+\tau_1\tau_2\tau_x-\tau_1\tau_2^2h_x}{\langle z_2\rangle}\Bigg)\,, \quad [\mathrm{S48}]$$

where

$$\psi = 2(\tau_2\tau_1^2h_x^2+\tau_2^2\tau_1h_x^2+\tau_2\tau_1^2+\tau_2^2\tau_1+\tau_1^2\tau_x+\tau_2\tau_1\tau_x+\tau_2^2\tau_x- 2\tau_2^2\tau_1h_x-2\tau_2\tau_1h_x\tau_x-\tau_2^2h_x\tau_x-2\tau_2\tau_1^2h_x-\tau_1^2h_x\tau_x)\,. \quad [\mathrm{S49}]$$

The stationary flux balance relations (see Eq. S7) for $Z_1$ and $Z_2$ ensures $\mu=\gamma\langle z_1z_2\rangle=\theta\langle x\rangle$ such that Eq. S2 implies $\tau_1=\langle z_1\rangle/(\theta\langle x\rangle)$ and $\tau_2=\langle z_2\rangle/(\theta\langle x\rangle)$ for this system. Substituting these lifetime identities and taking the limit of $\langle z_1\rangle\to\infty$ and $\langle z_2\rangle\to\infty$ of the approximate $\mathrm{CV}_x$ leads to (see Wolfram Notebook SupplementaryDerivations.nb)

$$\lim_{\langle z_1\rangle\to\infty}\lim_{\langle z_2\rangle\to\infty}\mathrm{CV}_x = \frac{1}{\sqrt{(1-h_x)\langle x\rangle}}\,. \quad [\mathrm{S50}]$$

Furthermore, we verify using the `Reduce[]` function in `Wolfram Mathematica 12` that this limit bounds all other physical solutions for the approximate $\mathrm{CV}_x$. Thus, we have the prediction

$$\mathrm{CV}_x \geq \frac{1}{\sqrt{(1-h_x)\langle x\rangle}} \,. \qquad \text{[S51]}$$

Thus, the linear noise approximation predicts noise suppression relative to an open-loop control strategy can be achieved by this closed-loop system, as expected (9). However, the right side of this inequality is exactly the linear noise approximation for $\mathrm{CV}_x$ for the system with only negative self-feedback with an equivalent non-linearity in the absence of the AIF input, i.e.,

$$\mathrm{CV}_x^{(\text{no AIF})} = \frac{1}{\sqrt{(1-h_x)\langle x\rangle}} \,, \qquad \text{[S52]}$$

see Wolfram Notebook SupplementaryDerivations.nb. Thus, Eq. S51 predicts the AIF module introduces a noise penalty relative to the system under only proportional control.

#### S2. Numerical simulation data and estimates of sampling error

**A. Evaluation of stationary first- and second-order moment invariants for the numerical simulation statistics.** At stationarity, the birth and death fluxes of any component in a Markovian process must balance. The flux balance equations for the general three-component AIF topology specified by Eq. S1 are given by Eq. S7. Analogously, Eq. 17 specifies a second-order moment invariant for pairs of components in a Markovian process (3, 7). These (co)variance balance relations for the general three-component AIF topology specified by Eq. S1 can be expressed as follows:

$$\begin{aligned}
\eta_{xx} &= \frac{\mathrm{Cov}(f(x,z_1,z_2),\ z)}{\langle f(x,z_1,z_2)\rangle\langle x\rangle} + \frac{1}{\langle x\rangle} \\
E_1\frac{\mathrm{Cov}(z_1z_2,\ z_2)}{\langle z_1z_2\rangle\langle z_2\rangle} + (1-E_1)\eta_{z_1z_1} &= \frac{1}{\langle z_1\rangle} \\
E_2\frac{\mathrm{Cov}(z_1z_2,\ z_2)}{\langle z_1z_2\rangle\langle z_2\rangle} + (1-E_2)\eta_{z_2z_2} &= \eta_{xz_2} + \frac{1}{\langle z_2\rangle} \\
\left(\frac{1-E_1}{\tau_1}+\frac{1-E_2}{\tau_2}\right)\eta_{z_1z_2} + \frac{E_1}{\tau_1}\frac{\mathrm{Cov}(z_1z_2,\ z_2)}{\langle z_1z_2\rangle\langle z_2\rangle} + \frac{E_2}{\tau_2}\frac{\mathrm{Cov}(z_1z_2,\ z_1)}{\langle z_1z_2\rangle\langle z_1\rangle} &= \\
\frac{1}{2}\left(\frac{E_1}{\tau_1\langle z_2\rangle}+\frac{E_2}{\tau_2\langle z_1\rangle}\right)& \\
\frac{1}{\tau_x}\eta_{xz_1} + \frac{E_1}{\tau_1}\eta_{xz_1} + \frac{1-E_1}{\tau_1}\frac{\mathrm{Cov}(z_1z_2,\ x)}{\langle z_1z_2\rangle\langle x\rangle} &= \frac{1}{\tau_x}\frac{\mathrm{Cov}(f(x,z_1,z_2),\ z_1)}{\langle f(x,z_1,z_2)\rangle\langle z_1\rangle} \\
\frac{1}{\tau_x}\eta_{xz_1} + \frac{E_1}{\tau_1}\eta_{xz_1} + \frac{1-E_1}{\tau_1}\frac{\mathrm{Cov}(z_1z_2,\ x)}{\langle z_1z_2\rangle\langle x\rangle} &= \frac{1}{\tau_x}\frac{\mathrm{Cov}(f(x,z_1,z_2),\ z_1)}{\langle f(x,z_1,z_2)\rangle\langle z_1\rangle}
\end{aligned} \qquad \text{[S53]}$$

Gillespie's stochastic simulation algorithm is an exact Monte Carlo procedure to generate sample trajectories for molecular abundances in any stochastic reaction network (10, 11). Thus, for a stable system it is a provably exact method for determining the stationary temporal joint distribution of molecular abundances in a stochastic reaction network in the limit of infinitely many simulation iterations. Practical considerations limit how many iterations the algorithm can be run for, which leads to inevitable sampling errors.

To avoid large sampling errors, we discard all simulation data that exhibit a relative error between the left and right side for the flux balance relations of Eq. S7 of more than 2%, or a relative error of the (co)variance balance relations of Eq. S53 of more than 5%. Typically, our observed numerical sampling errors in Eq. S7 and Eq. S53 are much smaller than the above cutoff, but *any* non-zero sampling error will inevitably lead to numerical errors even when checking exact analytical formulas. We thus evaluate the effect of the sampling error on our derived estimates by running at least three numerical simulations with distinct pseudo-random number generator seeds for any parameter set from initial parameter searches that violated our predictions derived from linear approximations. From these reseeded simulations, we calculate the mean and standard deviation of the numerical statistics to compare against our predictions to establish whether observed "violators" are statistically significant or due to numerical sampling error. Below this process is detailed for each system considered in this study.

**B. Idealized reference-actuated antithetic integral feedback system.** $1$ out of $1,716,668$ of the stationary $\mathrm{CV}_x$ values calculated in our numerical exploration (see Table S1) of the idealized reference-actuated AIF system fell below open-loop fluctuations given by $\mathrm{CV}_x = 1/\sqrt{\langle x \rangle}$, with a relative deviation of $\approx 0.0002$. However, when re-seeding simulations for this parameter set ten times for independent numerical realizations of the stochastic process, we obtained a mean $\mathrm{CV}_x$ that was $\approx 1.005$ times larger than the corresponding open-loop fluctuations. The corresponding standard deviations of the simulations was $\approx 0.0006$ suggesting that the initially observed "violator" was a statistical outlier due to inevitable sampling error in numerical simulations of systems close to the predicted bound.

For simulations satisfying $\mu/\theta \geq 1$ there were $20,789$ violators of the more restrictive lower bound of Eq. 6 predicted by the linear noise approximation (see Sect. S1.B for derivation) with a maximum relative deviation of $\approx 0.008$. Only 59 means across at least three seeds per violating parameter set fell more than one standard deviation from the bound predicted by Eq. 6, with a max relative deviation of the mean of $0.001$, corresponding to $\approx 1.1$ standard deviations. We ran this parameter set for several additional seeds for a total of 10 seeds, which resulted in the mean falling above the predicted bound (see Fig. S1A). This suggests that the small initial "violations" of the bound predicted by Eq. 6 were also due to inevitable sampling error in numerical simulations of systems close to the predicted bound.

In contrast, for simulations with $\mu/\theta < 1$ we found $116,096$ violators of the bound predicted by Eq. 6. Subsequent analysis suggest that many of them remain significant after re-sampling: We randomly selected the parameter sets for $4,024$ of these violators satisfying $\mu/\theta < 1$. The mean across at least three seeds fell more than one standard deviation below the predicted bound for $4,021$ of these parameter sets, with a maximum relative deviation of $\approx 0.03$, corresponding to $\approx 143$ standard deviations. Following the same process as for systems with $\mu/\theta \geq 1$, we ran this parameter set for several additional seeds for a total of 10 seeds. However, in contrast to our observations for $\mu/\theta \geq 1$, we observed negligible change in the mean and small standard deviations (see Fig. S1B). Although the violators were generally small (see Fig. S1B), this numerically suggests that Eq. 6 does not constrain systems with $\mu/\theta < 1$. In the main text we thus present the conclusion that numerical evidence suggests that Eq. 6 constrains only systems in which the average number of controlled molecules $X$ is at least one.

**C. Reference-actuated antithetic integral feedback system with dilution/degradation.** To compare our numerical simulations of the reference-actuated AIF system with dilution against the relationship of Eq. 10 predicted by our approximate derivation (see Sect. S1.D) we selected the parameter sets corresponding to the minimum $\mathrm{CV}_x$ from our numerical explorations (see Table S1) for each of

$N_2/N_x = 1,\ 5,\ 10,\ 20,\ 40,$ and 80 respectively. Note, the ratio $N_2/N_x$ is related to the simulation parameters through the relation $N_2/N_x = \theta\tau_x$ (see Sect. S1.B). We then ran ten independent replicate simulations for each parameter set, each with different pseudo-random number generator seeds. To evaluate the sampling error on the numerical $\mathrm{CV}_x^{\min}$ values and the corresponding sensitivity values $S^*$, we then calculated the mean and standard deviation of the numerical $\mathrm{CV}_x$ and numerical sensitivity coefficient across the six seeds for each $N_2/N_x$ value. The max relative deviation in both the $S^*$ and $\mathrm{CV}_x^{\min}$ dimensions of the prediction was exhibited for $N_2/N_x = 80$. In the $S^*$ dimension, the max relative deviation was $\approx 0.14$, corresponding to $\approx 17$ standard deviations. The max relative deviation in the $\mathrm{CV}_x^{\min}$ dimension of the prediction was $\approx 0.039$ for $N_2/N_x = 80$, corresponding to $\approx 56$ standard deviations. The max relative deviation from the approximate theory in both the $S^*$ and $\mathrm{CV}_x^{\min}$ dimensions is small relative to the predicted value, but the small standard deviations relative to these deviations from the approximate theory suggests the deviation is not explained by sampling error. This suggests that that the prediction of Eq. 10, which is only realized in the optimal limit, is not exactly realized by any of our parameter sets, but they do come very close. We thus conclude that the numerical simulations suggest that the approximate theory accurately predicts the maximum noise suppression and corresponding sensitivity for a given $N_2/N_x$. Note, we also evaluated the numerical sampling error on the data presented in Fig. 3 (see Fig. S5).

**D. Idealized sensor-actuated antithetic integral feedback system.** To confirm that numerical $\mathrm{CV}_x$ values below open-loop levels for the idealized sensor actuated system were not merely the result of numerical sampling error, we followed the same procedure used in Sect. S2.B to reject potential outliers for the idealized reference-actuated system.

$11,169$ out of the $27,520$ stationary $\mathrm{CV}_x$ values calculated in our numerical exploration (see Table S1) of the idealized sensor-actuated AIF system fell below open-loop fluctuations given by $\mathrm{CV}_x = 1/\sqrt{\langle x \rangle}$, with a max relative deviation below open-loop fluctuations of $\approx 0.45$. For each of the parameter sets corresponding to the ten smallest values of $\mathrm{CV}_x/(1/\sqrt{x}))$ observed in our numerical exploration (see Table S1), we ran five numerical simulations, each for a different pseudo-random number generator seed. The mean $\mathrm{CV}_x$ across the five seeds fell at least one standard deviation below $1/\sqrt{\langle x \rangle}$ for each considered parameter set. The *minimum* relative deviation below open-loop levels amongst these reseeded parameter sets was $\approx 0.42$, corresponding to $\approx 4900$ standard deviations. We ran this parameter set for several additional seeds for a total of 10 seeds, which showed negligible change in the mean and small standard deviations (see Fig. S2A). This confirms that sensor-actuated AIF can achieve noise suppression below uncontrolled levels. This conclusion is further supported by a reseeding analysis of a second parameter set showing even greater noise suppression despite more biologically realistic parameters (see Fig. S2B).

357 # S3. Supplementary figures and table

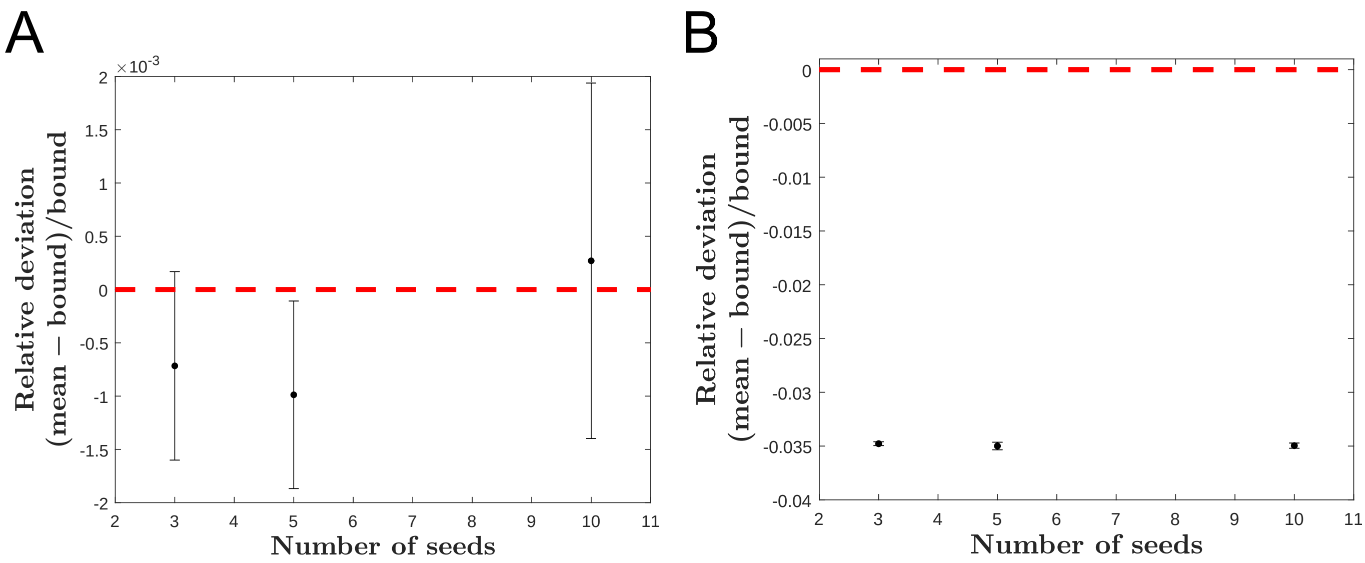

**Fig. S1. Evaluating sampling error for most severe violations of Eq. 6.** A) Example data for the most severe (but still minuscule) outlier below the bound predicted by Eq. 6 in a system with $\mu/\theta \geq 1$. The data set was resampled independently for a total of ten unique pseudo-random number generator seeds. The plot depicts the dependence of the relative deviation on the number of seeds used to compute the mean $\mathrm{CV}_x$ for this parameter set. Error bars are given as the standard deviation of the $\mathrm{CV}_x$ across the seeds. Eventually, when ten seeds are considered, the mean falls above the bound predicted by Eq. 6. Note also, the standard deviation is larger when computed across ten seeds, compared to for three and five seeds, indicating significant sampling error which is underestimated for smaller seed numbers. Note, systems featuring complex formation motifs are known to exhibit multimodal behaviour (3), which may explain the large sampling error on this point and others considered in our reseeding procedure. B) Here, the same procedure was followed as in A) for the most severe outlier below the bound predicted by Eq. 6 from the reseeding procedure satisfying $\mu/\theta < 1$. In contrast to A) We observe very little change in the mean $\mathrm{CV}_x$ for an increasing number of seeds as well as a negligible standard deviation compared to the relative deviation below the bound predicted by Eq. 6. This suggests Eq. 6 does not constrain systems with $\mu/\theta < 1$.

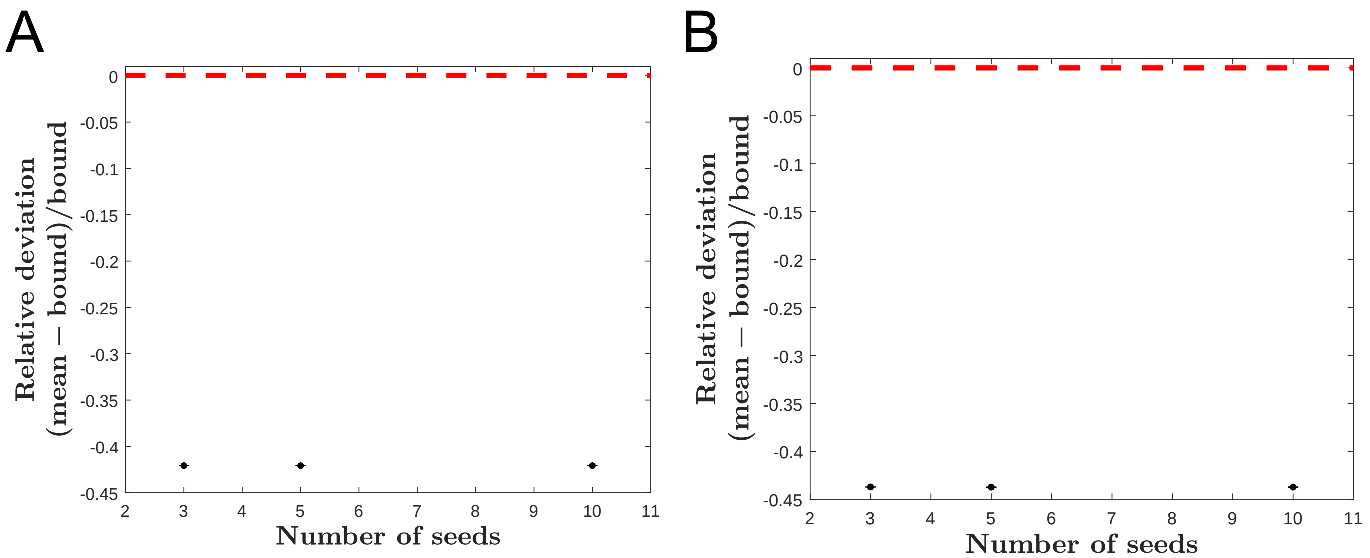

**Fig. S2. Evaluating sampling error for idealized sensor-actuated systems exhibiting noise suppression.** A) Ten parameter sets corresponding to the most extreme noise suppression values observed in the initial parameter search for the sensor-actuated system were simulated for five unique pseudo-random number generator seeds each. The parameter set whose mean $\mathrm{CV}_x$ from the reseeding procedure exhibited the smallest relative deviation below open-loop fluctuations was reseeded several additional times for a total of ten unique pseudo-random number generator seeds. The plot depicts the dependence of the relative deviation on the number of seeds used to compute the mean $\mathrm{CV}_x$ for this parameter set. Error bars are given as the standard deviation of the $\mathrm{CV}_x$ across the seeds. We observe very little change in the mean $\mathrm{CV}_x$ for an increasing number of seeds as well as a negligible standard deviation compared to the relative deviation below open-loop fluctuations. This suggests the points exhibiting noise suppression are *not* merely the result of sampling error. B) The absolute Hill coefficient for the parameter set considered in A) was $\approx 30$, which is biologically implausible. Here, the same procedure was followed as in A) for a parameter set with a more realistic absolute Hill coefficient of $\approx 4$. Again, we observe very little change in the mean $\mathrm{CV}_x$ for an increasing number of seeds as well as a negligible standard deviation compared to the relative deviation below open-loop fluctuations. Moreover, we see that this parameter set with a more realistic Hill coefficient exhibits even greater noise suppression than the parameter set considered in A).

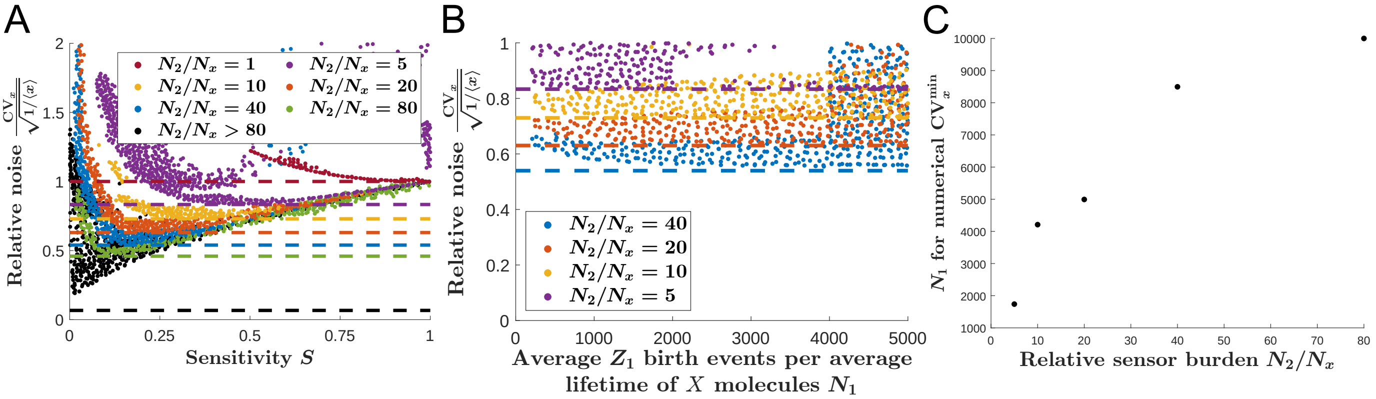

**Fig. S3. Supplementary data for reference-actuated antithetic integral feedback system with dilution.** A) Shown are noise levels and sensitivity coefficients $S$ as defined in Eq. 8 for an illustrative sub-sample of numerical simulations with different control parameters (see Materials and Methods; see also Table S1). Data points for the numerical simulations are coloured according the value of $N_2/N_x$. The noise floor predicted by Eq. 9 is plotted as a dashed line for each $N_2/N_x$ value with the same colour as the dots it is predicted to constrain. The black dashed line corresponds to the noise floor predicted by Eq. 9 for $N_2/N_x = 200,000$, which is the largest $N_2/N_x$ value sampled by our numerical exploration. Note, due to the numerical simulations becoming exceedingly expensive computationally, we do not sample near the predicted noise floor in the $N_2/N_x > 80$ regime. However, for each $N_2/N_x < 80$ value, we see Eq. 9 accurately predicts the minimum attainable relative noise. While our numerical data establishes there is no fundamental trade-off between sensitivity and noise, it does suggests that for a fixed $N_2/N_x$ value fluctuations must be made larger to decrease sensitivity below $1/\sqrt{N_2/N_x}$, where the noise floor is attainable (see Sect. S1.D). B) Shown are noise levels and the average number of $Z_1$ birth events per average controlled species lifetime $\tau_x = 1/\beta_x$, given by $N_1$. We plot an illustrative sub-sample of numerical simulations with different control parameters (see Materials and Methods; see also Table S1) for $N_2/N_x = 5,\ 10,\ 20,\ 40$ to illustrate that our data suggests the value of $N_1$ restricts how close the noise can get to the noise floor predicted by Eq. 9. Data points for the numerical simulations are coloured according the value of $N_2/N_x$, and the corresponding noise floor predicted by Eq.. 9 is given a dashed line of the same colour. C) Here we plot the $N_1$ value corresponding the minimum relative noise values observed in the numerical exploration used to generate Fig. 3C against the relative sensor burden $N_2/N_x$. We observe that larger $N_2/N_x$ values corresponded to larger $N_1$ values for the numerical simulation data to approach the noise floor predicted by Eq. 9. This is why probing the predicted noise floor for $N_2/N_x > 80$ becomes exceedingly computationally expensive.

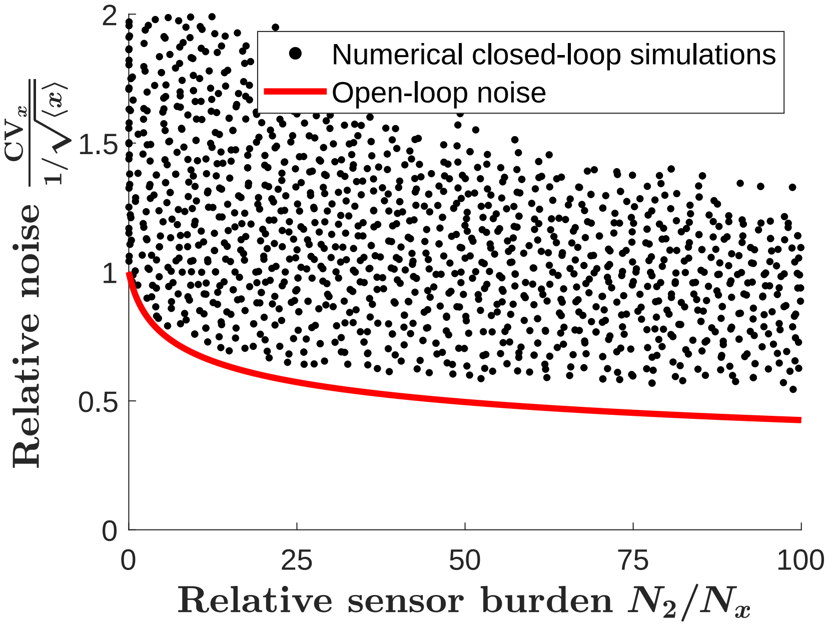

**Fig. S4. Comparison of numerical simulations of the idealized sensor-actuated AIF system with the bound predicted by Eq. 13.** Eq. 13 predicts that noise suppression in our minimal model of sensor-actuated AIF control (Fig. 4A; see also Eq. 11) is bounded from below by a decreasing function that scales with the quartic root of the relative control burden $2N_2/N_x$. Numerical simulations (black dots) satisfy this prediction. Shown are a representative sub-sample of a parameter search for such noise suppressing systems that achieve perfect adaptation (see Table S1). This suggests arbitrary noise suppression and perfect adaption can be achieve by idealized sensor-actuated AIF so long as cells can afford to pay the corresponding relative control burden given by $2N_2/N_x$.

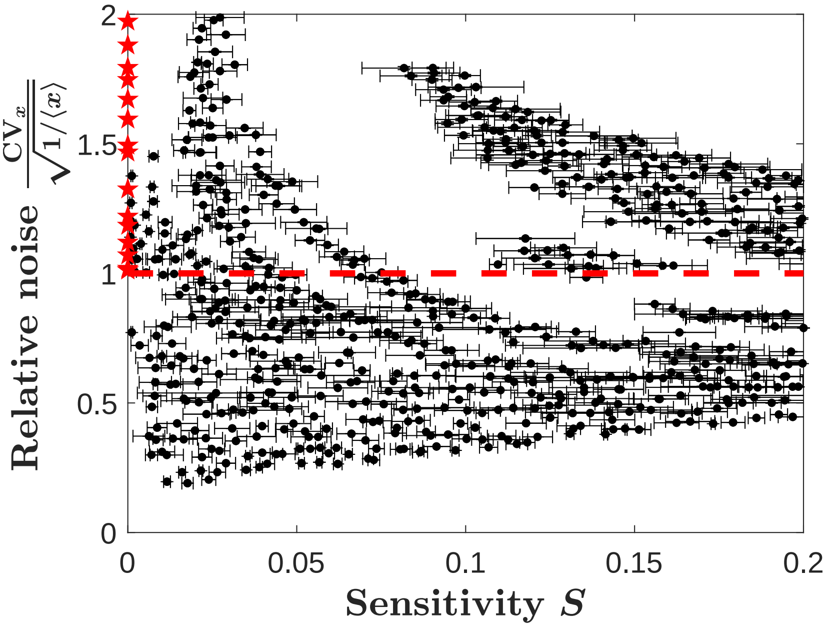

**Fig. S5. Evaluating sampling error for reference-actuated with dilution/degradation.** To address inevitable sampling error when numerically estimating sensitivity and noise in the reference-actuated system with dilution/degradation, we simulated each parameter set independently for at least three unique pseudo-random number generator seeds. The data points plotted in Fig. 3 correspond to the mean sensitivity and noise values across the seeds. Here we plot this same data along with the corresponding standard deviation values calculated across the seeds. We observe sampling error has a more significant effect on the numerical sensitivity values than the corresponding relative noise values. However, taking this variation in the sensitivity estimates into account does not significantly change the interpretation of the plot: significant noise suppression is exhibited for marginal sensitivity values.

**Table S1. Parameter sampling for numerical simulations.** Without loss of generality, all simulations of idealized systems were performed with $\beta_x \equiv 1$ such that all rates can be regarded in units of $\beta_x$. For systems with control species dilution ($\beta_1,\ \beta_2 \neq 0$), two simulations per parameter set were run, one with $\beta_x = 0.99$ and one with $\beta_x = 1.01$, to compute the numerical sensitivity coefficient (see Eq. 23). The control input function for idealized sensor-actuated AIF simulations was taken to be the Hill function $f(z_2) = \nu z_2^n/(\kappa^n + z_2^n)$.

| System | Sampling range | Sampling distribution | Number of samples | Number satisfying flux & covariance balance test |
|---|---|---|---|---|
| Idealized reference-actuated AIF | $\mu$ : $0.1 \to 10$<br>$\theta$ : $0.1 \to 10$<br>$k$ : $0.1 \to 10$<br>$\gamma$ : $0.1 \to 10$ | logarithmic<br>logarithmic<br>logarithmic<br>logarithmic | $1,048,576$ | $803,331$ |
| | $\mu$ : $0.01 \to 100$<br>$\theta$ : $0.01 \to 100$<br>$k$ : $0.01 \to 100$<br>$\gamma$ : $0.01 \to 100$ | logarithmic<br>logarithmic<br>logarithmic<br>logarithmic | $1,424,008$ | $821,880$ |
| | $\mu$ : $0.1 \to 10$<br>$\theta = \mu$<br>$k$ : $0.1 \to 10$<br>$\gamma$ : $0.1 \to 10$ | logarithmic<br>logarithmic<br>logarithmic<br>logarithmic | $125,000$ | $91,457$ |
| Reference-actuated AIF with dilution/degradation | $\mu = 5,000$<br>$\theta = 200,000$<br>$k = 0.6667$<br>$\gamma = 1,350.95$<br>$\beta_1 = 100$<br>$\beta_2 = 100$ | N/A<br>N/A<br>N/A<br>N/A<br>N/A<br>N/A | 1 | 1 |
| | $\mu$ : $200 \to 5,000$<br>$\theta = 1$<br>$k$ : $10 \to 100$<br>$\gamma$ : $500 \to 5,000$<br>$\beta_1 = 100$<br>$\beta_2 = 100$ | uniform<br>N/A<br>uniform<br>uniform<br>N/A<br>N/A | 1,974 | 1,974 |
| | $\mu$ : $200 \to 5,000$<br>$\theta = 5$<br>$k$ : $200 \to 1,000$<br>$\gamma$ : $500 \to 5,000$<br>$\beta_1 = 100$<br>$\beta_2 = 100$ | uniform<br>N/A<br>uniform<br>uniform<br>N/A<br>N/A | 5,000 | 4,441 |
| | $\mu$ : $200 \to 5,000$<br>$\theta = 5$<br>$k$ : $200 \to 1,000$<br>$\gamma$ : $0 \to 500$<br>$\beta_1 = 100$<br>$\beta_2 = 100$ | uniform<br>N/A<br>uniform<br>uniform<br>N/A<br>N/A | 5,000 | 2,770 |

| | | | | |
|---|---|---|---|---|
| | $\mu\ :\ 200 \to 5,000$<br>$\theta = 5$<br>$k\ :\ 200 \to 1,000$<br>$\gamma\ :\ 0 \to 1$<br>$\beta_1 = 100$<br>$\beta_2 = 100$ | uniform<br>N/A<br>uniform<br>uniform<br>N/A<br>N/A | 5,000 | 392 |
| | $\mu\ :\ 200 \to 5,000$<br>$\theta = 5$<br>$k\ :\ 20 \to 30$<br>$\gamma = 300$<br>$\beta_1 = 1000$<br>$\beta_2 = 1000$ | uniform<br>N/A<br>uniform<br>N/A<br>N/A<br>N/A | 1,262 | 837 |
| | $\mu\ :\ 4,000 \to 5,000$<br>$\theta = 5$<br>$k\ :\ 20 \to 30$<br>$\gamma = 300$<br>$\beta_1 = 100$<br>$\beta_2 = 100$ | uniform<br>N/A<br>uniform<br>N/A<br>N/A<br>N/A | 5,000 | 4,995 |
| | $\mu\ :\ 200 \to 5,000$<br>$\theta = 10$<br>$k\ :\ 10 \to 100$<br>$\gamma\ :\ 500 \to 5,000$<br>$\beta_1 = 100$<br>$\beta_2 = 100$ | uniform<br>N/A<br>uniform<br>uniform<br>N/A<br>N/A | 5,000 | 4,000 |
| | $\mu\ :\ 200 \to 5,000$<br>$\theta = 20$<br>$k\ :\ 10 \to 100$<br>$\gamma\ :\ 500 \to 5,000$<br>$\beta_1 = 100$<br>$\beta_2 = 100$ | uniform<br>N/A<br>uniform<br>uniform<br>N/A<br>N/A | 5,000 | 4,137 |
| | $\mu\ :\ 4,000 \to 5,000$<br>$\theta = 20$<br>$k\ :\ 20 \to 30$<br>$\gamma = 300$<br>$\beta_1 = 100$<br>$\beta_2 = 100$ | uniform<br>N/A<br>uniform<br>N/A<br>N/A<br>N/A | 5,000 | 2,605 |
| | $\mu\ :\ 4,000 \to 5,000$<br>$\theta = 20$<br>$k\ :\ 100 \to 200$<br>$\gamma = 300$<br>$\beta_1 = 100$<br>$\beta_2 = 100$ | uniform<br>N/A<br>uniform<br>N/A<br>N/A<br>N/A | 5,000 | 1,985 |
| | $\mu\ :\ 6,000 \to 7,000$<br>$\theta = 20$<br>$k\ :\ 300 \to 400$<br>$\gamma = 300$<br>$\beta_1 = 100$<br>$\beta_2 = 100$ | uniform<br>N/A<br>uniform<br>N/A<br>N/A<br>N/A | 5,000 | 1,120 |

| | | | | |
|---|---|---|---|---|
| | $\mu\ :\ 4,000 \to 5,000$<br>$\theta = 20$<br>$k\ :\ 100 \to 200$<br>$\gamma = 300$<br>$\beta_1 = 100$<br>$\beta_2 = 100$ | uniform<br>N/A<br>uniform<br>N/A<br>N/A<br>N/A | 5,000 | 1,982 |
| | $\mu\ :\ 4,000 \to 5,000$<br>$\theta = 20$<br>$k\ :\ 100 \to 200$<br>$\gamma = 300$<br>$\beta_1\ :\ 0 \to 1,000$<br>$\beta_2 = \beta_1$ | uniform<br>N/A<br>uniform<br>N/A<br>uniform<br>N/A | 5,000 | 587 |
| | $\mu\ :\ 200 \to 5,000$<br>$\theta = 40$<br>$k\ :\ 10 \to 100$<br>$\gamma\ :\ 500 \to 4,500$<br>$\beta_1 = 100$<br>$\beta_2 = 100$ | uniform<br>N/A<br>uniform<br>uniform<br>N/A<br>N/A | 5,000 | 4831 |
| | $\mu\ :\ 200 \to 6,500$<br>$\theta = 40$<br>$k\ :\ 200 \to 2,0000$<br>$\gamma\ :\ 0 \to 1$<br>$\beta_1 = 100$<br>$\beta_2 = 100$ | uniform<br>N/A<br>uniform<br>uniform<br>N/A<br>N/A | 5,000 | 245 |
| | $\mu\ :\ 500 \to 1,000$<br>$\theta = 40$<br>$k\ :\ 10 \to 100$<br>$\gamma = 300$<br>$\beta_1 = 100$<br>$\beta_2 = 100$ | uniform<br>N/A<br>uniform<br>N/A<br>N/A<br>N/A | 5,000 | 4,932 |
| | $\mu\ :\ 4,000 \to 5,000$<br>$\theta = 40$<br>$k\ :\ 20 \to 30$<br>$\gamma = 300$<br>$\beta_1 = 100$<br>$\beta_2 = 100$ | uniform<br>N/A<br>uniform<br>N/A<br>N/A<br>N/A | 5,000 | 4,601 |
| | $\mu\ :\ 4,000 \to 5,000$<br>$\theta = 40$<br>$k\ :\ 100 \to 200$<br>$\gamma = 300$<br>$\beta_1\ :\ 0 \to 100$<br>$\beta_2 = \beta_1$ | uniform<br>N/A<br>uniform<br>N/A<br>uniform<br>N/A | 5,000 | 4,545 |
| | $\mu\ :\ 4,000 \to 5,000$<br>$\theta = 80$<br>$k\ :\ 100 \to 200$<br>$\gamma = 300$<br>$\beta_1 = 100$<br>$\beta_2 = 100$ | uniform<br>N/A<br>uniform<br>N/A<br>N/A<br>N/A | 5,000 | 3,266 |

| | | | | |
|---|---|---|---|---|
| | $\mu\ :\ 4,000 \to 5,000$<br>$\theta = 80$<br>$k\ :\ 0 \to 1$<br>$\gamma = 300$<br>$\beta_1 = 100$<br>$\beta_2 = 100$ | uniform<br>N/A<br>uniform<br>N/A<br>N/A<br>N/A | 4,999 | 2,899 |
| | $\mu\ :\ 4,000 \to 5,000$<br>$\theta = 80$<br>$k\ :\ 1 \to 100$<br>$\gamma = 300$<br>$\beta_1 = 100$<br>$\beta_2 = 100$ | uniform<br>N/A<br>uniform<br>N/A<br>N/A<br>N/A | 5,000 | 4,342 |
| | $\mu\ :\ 5,000 \to 6,000$<br>$\theta = 80$<br>$k = 10$<br>$\gamma\ :\ 200 \to 400$<br>$\beta_1 = 100$<br>$\beta_2 = 100$ | uniform<br>N/A<br>N/A<br>uniform<br>N/A<br>N/A | 5,000 | 4,584 |
| | $\mu\ :\ 400 \to 4,000$<br>$\theta\ :\ 13 \to 134$<br>$k\ :\ 2 \to 20$<br>$\gamma\ :\ 800 \to 8,000$<br>$\beta_1\ :\ 20 \to 200$<br>$\beta_2 = \beta_1$ | uniform<br>uniform<br>uniform<br>uniform<br>uniform<br>N/A | 5,000 | 5,000 |
| | $\mu\ :\ 400 \to 4,000$<br>$\theta = \mu/30$<br>$k\ :\ 2 \to 20$<br>$\gamma\ :\ 800 \to 8,000$<br>$\beta_1\ :\ 1,000 \to 10,000$<br>$\beta_2 = \beta_1$ | uniform<br>N/A<br>uniform<br>uniform<br>uniform<br>N/A | 1,992 | 1,935 |
| | $\mu\ :\ 200 \to 1,000$<br>$\theta = 5$<br>$k\ :\ 20 \to 80$<br>$\gamma\ :\ 500 \to 5,000$<br>$\beta_1 = 100$<br>$\beta_2 = 100$ | uniform<br>N/A<br>uniform<br>uniform<br>N/A<br>N/A | 5,000 | 5,000 |
| | $\mu\ :\ 200 \to 1,500$<br>$\theta = 100$<br>$k\ :\ 10 \to 50$<br>$\gamma\ :\ 400 \to 1,000$<br>$\beta_1 = 100$<br>$\beta_2 = 100$ | uniform<br>N/A<br>uniform<br>uniform<br>N/A<br>N/A | 5,000 | 4,998 |
| | $\mu\ :\ 1,000 \to 5,000$<br>$\theta = 100$<br>$k\ :\ 10 \to 50$<br>$\gamma\ :\ 400 \to 1,000$<br>$\beta_1 = 100$<br>$\beta_2 = 100$ | uniform<br>N/A<br>uniform<br>uniform<br>N/A<br>N/A | 5,000 | 5,000 |

| | | | | |
|---|---|---|---|---|
| | $\mu\ :\ 6,000 \to 10,000$<br>$\theta = 100$<br>$k\ :\ 10 \to 50$<br>$\gamma\ :\ 400 \to 1,000$<br>$\beta_1 = 100$<br>$\beta_2 = 100$ | uniform<br>N/A<br>uniform<br>uniform<br>N/A<br>N/A | 5,000 | 4,120 |
| | $\mu\ :\ 1,000 \to 2,0000$<br>$\theta = 100$<br>$k\ :\ 0 \to 200$<br>$\gamma\ :\ 500 \to 5,000$<br>$\beta_1 = 100$<br>$\beta_2 = 100$ | uniform<br>N/A<br>uniform<br>uniform<br>N/A<br>N/A | 5,000 | 4,985 |
| | $\mu\ :\ 1,000 \to 2,0000$<br>$\theta = 5$<br>$k\ :\ 0 \to 200$<br>$\gamma\ :\ 500 \to 5,000$<br>$\beta_1 = 100$<br>$\beta_2 = 100$ | uniform<br>N/A<br>uniform<br>uniform<br>N/A<br>N/A | 4,999 | 4,807 |
| | $\mu\ :\ 7,000 \to 10,000$<br>$\theta = 80$<br>$k\ :\ 1 \to 100$<br>$\gamma = 300$<br>$\beta_1 = 100$<br>$\beta_2 = 100$ | uniform<br>N/A<br>logarithmic<br>N/A<br>N/A<br>N/A | 453 | 424 |
| | $\mu = 10,000$<br>$\theta = 80$<br>$k\ :\ 1 \to 100$<br>$\gamma = 300$<br>$\beta_1\ :\ 10 \to 100$<br>$\beta_2 = \beta_1$ | N/A<br>N/A<br>logarithmic<br>N/A<br>logarithmic<br>N/A | 500 | 397 |
| | $\mu\ :\ 2,500 \to 5,000$<br>$\theta\ :\ 100 \to 500$<br>$k\ :\ 0.05 \to 5$<br>$\gamma\ :\ 500 \to 5,000$<br>$\beta_1\ :\ 0.1 \to 1$<br>$\beta_2 = \beta_1$ | uniform<br>uniform<br>logarithmic<br>logarithmic<br>logarithmic<br>N/A | 4,429 | 3,890 |
| | $\mu\ :\ 2,500 \to 5,000$<br>$\theta\ :\ 500 \to 1,00$<br>$k\ :\ 0.05 \to 5$<br>$\gamma\ :\ 500 \to 5,000$<br>$\beta_1\ :\ 0.1 \to 1$<br>$\beta_2 = \beta_1$ | uniform<br>uniform<br>logarithmic<br>logarithmic<br>logarithmic<br>N/A | 1,312 | 993 |
| | $\mu\ :\ 2,500 \to 5,000$<br>$\theta\ :\ 100 \to 500$<br>$k\ :\ 0.05 \to 5$<br>$\gamma\ :\ 500 \to 5,000$<br>$\beta_1\ :\ 0.01 \to 0.1$<br>$\beta_2 = \beta_1$ | uniform<br>uniform<br>logarithmic<br>logarithmic<br>logarithmic<br>N/A | 426 | 423 |

| | | | | |
|---|---|---|---|---|
| | $\mu\ :\ 200 \to 500$<br>$\theta\ :\ 100 \to 500$<br>$k\ :\ 0.01 \to 10$<br>$\gamma\ :\ 100 \to 500$<br>$\beta_1 = 100$<br>$\beta_2 = 100$ | uniform<br>uniform<br>logarithmic<br>uniform<br>N/A<br>N/A | 715 | 715 |
| | $\mu\ :\ 2,500 \to 5,000$<br>$\theta\ :\ 500 \to 1,000$<br>$k = 0.1$<br>$\gamma = 500$<br>$\beta_1\ :\ 0.01 \to 0.1$<br>$\beta_2 = \beta_1$ | uniform<br>uniform<br>N/A<br>N/A<br>logarithmic<br>N/A | 500 | 369 |
| | $\mu\ :\ 2,500 \to 5,000$<br>$\theta\ :\ 500 \to 1,000$<br>$k = 0.1$<br>$\gamma = 500$<br>$\beta_1\ :\ 0.1 \to 1$<br>$\beta_2 = \beta_1$ | uniform<br>uniform<br>N/A<br>N/A<br>logarithmic<br>N/A | 500 | 331 |
| | $\mu = 5,000$<br>$\theta = 100$<br>$k\ :\ 0.05 \to 0.5$<br>$\gamma = 500$<br>$\beta_1\ :\ 0.1 \to 1$<br>$\beta_2 = \beta_1$ | N/A<br>N/A<br>logarithmic<br>N/A<br>logarithmic<br>N/A | 45 | 44 |
| | $\mu = 5,000$<br>$\theta = 100$<br>$k\ :\ 0.05 \to 0.5$<br>$\gamma = 500$<br>$\beta_1\ :\ 0.01 \to 0.1$<br>$\beta_2 = \beta_1$ | N/A<br>N/A<br>logarithmic<br>N/A<br>logarithmic<br>N/A | 44 | 43 |
| | $\mu = 5,000$<br>$\theta = 100$<br>$k\ :\ 0.05 \to 0.5$<br>$\gamma = 500$<br>$\beta_1\ :\ 1 \to 10$<br>$\beta_2 = \beta_1$ | N/A<br>N/A<br>logarithmic<br>N/A<br>logarithmic<br>N/A | 95 | 94 |
| | $\mu = 5,000$<br>$\theta = 100$<br>$k\ :\ 0.05 \to 0.5$<br>$\gamma = 500$<br>$\beta_1\ :\ 10 \to 100$<br>$\beta_2 = \beta_1$ | N/A<br>N/A<br>logarithmic<br>N/A<br>logarithmic<br>N/A | 100 | 96 |
| | $\mu\ :\ 1,000 \to 10,000$<br>$\theta\ :\ 1,000 \to 10,000$<br>$k = 0.4$<br>$\gamma = 500$<br>$\beta_1 = 1,000$<br>$\beta_2 = 1,000$ | logarithmic<br>logarithmic<br>N/A<br>N/A<br>N/A<br>N/A | 100 | 71 |

| | | | | |
|---|---|---|---|---|
| | $\mu$ : $1,000 \to 10,000$<br>$\theta$ : $1,000 \to 10,000$<br>$k = 0.4$<br>$\gamma$ : $1,000 \to 10,000$<br>$\beta_1 = 1,000$<br>$\beta_2 = 1,000$ | logarithmic<br>logarithmic<br>N/A<br>logarithmic<br>N/A<br>N/A | 100 | 78 |
| | $\mu$ : $50 \to 500,000$<br>$\theta = 10,000$<br>$k$ : $0.5 \to 500$<br>$\gamma$ : $10,000 \to 100,000$<br>$\beta_1 = 1,000$<br>$\beta_2 = 1,000$ | logarithmic<br>N/A<br>logarithmic<br>logarithmic<br>N/A<br>N/A | 50 | 42 |
| | $\mu$ : $60 \to 60,000$<br>$\theta = 1,000$<br>$k$ : $2 \to 20$<br>$\gamma$ : $10 \to 1,00$<br>$\beta_1 = 100$<br>$\beta_2 = 100$ | logarithmic<br>N/A<br>logarithmic<br>logarithmic<br>N/A<br>N/A | 50 | 33 |
| | $\mu$ : $30 \to 30,000$<br>$\theta = 100$<br>$k$ : $15 \to 150$<br>$\gamma$ : $200 \to 2,000$<br>$\beta_1 = 100$<br>$\beta_2 = 100$ | logarithmic<br>N/A<br>logarithmic<br>logarithmic<br>N/A<br>N/A | 50 | 42 |
| | $\mu$ : $30,000 \to 300,000$<br>$\theta = 100$<br>$k$ : $15 \to 150$<br>$\gamma$ : $200 \to 2,000$<br>$\beta_1 = 100$<br>$\beta_2 = 100$ | logarithmic<br>N/A<br>logarithmic<br>logarithmic<br>N/A<br>N/A | 50 | 11 |
| | $\mu$ : $50 \to 500,000$<br>$\theta = 10,000$<br>$k$ : $0.5 \to 500$<br>$\gamma$ : $10,000 \to 100,000$<br>$\beta_1 = 10,000$<br>$\beta_2 = 10,000$ | logarithmic<br>N/A<br>logarithmic<br>logarithmic<br>N/A<br>N/A | 50 | 32 |
| | $\mu$ : $50 \to 500,000$<br>$\theta = 10,000$<br>$k$ : $0.5 \to 500$<br>$\gamma$ : $10,000 \to 100,000$<br>$\beta_1 = 100$<br>$\beta_2 = 100$ | logarithmic<br>N/A<br>logarithmic<br>logarithmic<br>N/A<br>N/A | 440 | 426 |
| | $\mu$ : $50 \to 500,000$<br>$\theta = 10,000$<br>$k$ : $0.5 \to 500$<br>$\gamma$ : $10,000 \to 100,000$<br>$\beta_1 = 10$<br>$\beta_2 = 10$ | logarithmic<br>N/A<br>logarithmic<br>logarithmic<br>N/A<br>N/A | 40 | 38 |

| | | | | |
|---|---|---|---|---|
| | $\mu$ : $50 \to 500,000$<br>$\theta$ : $1,000 \to 10,000$<br>$k$ : $0.5 \to 500$<br>$\gamma$ : $200 \to 200,000$<br>$\beta_1$ : $10 \to 100$<br>$\beta_2$ : $10 \to 100$ | logarithmic<br>logarithmic<br>logarithmic<br>logarithmic<br>logarithmic<br>logarithmic | 100 | 95 |
| | $\mu$ : $50 \to 500,000$<br>$\theta$ : $500 \to 5,000$<br>$k$ : $0.5 \to 500$<br>$\gamma$ : $200 \to 200,000$<br>$\beta_1$ : $10 \to 100$<br>$\beta_2$ : $10 \to 100$ | logarithmic<br>logarithmic<br>logarithmic<br>logarithmic<br>logarithmic<br>logarithmic | 302 | 276 |
| | $\mu$ : $50 \to 500,000$<br>$\theta$ : $500 \to 5,000$<br>$k$ : $0.5 \to 500$<br>$\gamma$ : $200 \to 200,000$<br>$\beta_1$ : $1 \to 10$<br>$\beta_2$ : $1 \to 10$ | logarithmic<br>logarithmic<br>logarithmic<br>logarithmic<br>logarithmic<br>logarithmic | 498 | 425 |
| | $\mu$ : $50 \to 50,000$<br>$\theta$ : $500 \to 5,000$<br>$k$ : $0.5 \to 500$<br>$\gamma$ : $200 \to 200,000$<br>$\beta_1$ : $10 \to 100$<br>$\beta_2$ : $10 \to 100$ | logarithmic<br>logarithmic<br>logarithmic<br>logarithmic<br>logarithmic<br>logarithmic | 80 | 68 |
| | $\mu$ : $50 \to 50,000$<br>$\theta$ : $500 \to 5,000$<br>$k$ : $0.5 \to 500$<br>$\gamma$ : $200 \to 200,000$<br>$\beta_1$ : $10 \to 100$<br>$\beta_2$ : $10 \to 100$ | logarithmic<br>logarithmic<br>logarithmic<br>logarithmic<br>logarithmic<br>logarithmic | 500 | 478 |
| | $\mu = 455.858595$<br>$\theta = 1$<br>$k = 0.691554$<br>$\gamma = 103.681817$<br>$\beta_1 = 100$<br>$\beta_2 = 100$ | N/A<br>N/A<br>N/A<br>N/A<br>N/A<br>N/A | 15 | 15 |
| | $\mu = 1738.603162$<br>$\theta = 5$<br>$k = 24.387861$<br>$\gamma = 4,722.311054$<br>$\beta_1 = 100$<br>$\beta_2 = 100$ | N/A<br>N/A<br>N/A<br>N/A<br>N/A<br>N/A | 15 | 15 |
| | $\mu = 4210.045673$<br>$\theta = 10$<br>$k = 23.897082$<br>$\gamma = 4,720.616247$<br>$\beta_1 = 100$<br>$\beta_2 = 100$ | N/A<br>N/A<br>N/A<br>N/A<br>N/A<br>N/A | 15 | 15 |

| | | | | |
|---|---|---|---|---|
| | $\mu = 4,992.856532$<br>$\theta = 20$<br>$k = 19.986089$<br>$\gamma = 1,451.220664$<br>$\beta_1 = 100$<br>$\beta_2 = 100$ | N/A<br>N/A<br>N/A<br>N/A<br>N/A<br>N/A | 15 | 15 |
| | $\mu = 4,809.170535$<br>$\theta = 40$<br>$k = 15.184838$<br>$\gamma = 4,227.122626$<br>$\beta_1 = 100$<br>$\beta_2 = 100$ | N/A<br>N/A<br>N/A<br>N/A<br>N/A<br>N/A | 20 | 20 |
| | $\mu = 10,000$<br>$\theta = 80$<br>$k = 10.478416$<br>$\gamma = 300$<br>$\beta_1 = 79.152995$<br>$\beta_2 = 79.152995$ | N/A<br>N/A<br>N/A<br>N/A<br>N/A<br>N/A | 20 | 20 |
| | $\mu = 4,558.585954$<br>$\theta = 1$<br>$k = 0.691554$<br>$\gamma = 103.681817$<br>$\beta_1 = 100$<br>$\beta_2 = 100$ | N/A<br>N/A<br>N/A<br>N/A<br>N/A<br>N/A | 1 | 1 |
| | $\mu = 17,386.031616$<br>$\theta = 5$<br>$k = 24.387861$<br>$\gamma = 4,722.311054$<br>$\beta_1 = 100$<br>$\beta_2 = 100$ | N/A<br>N/A<br>N/A<br>N/A<br>N/A<br>N/A | 1 | 1 |
| | $\mu = 42,100.456732$<br>$\theta = 10$<br>$k = 23.897082$<br>$\gamma = 4,720.616247$<br>$\beta_1 = 100$<br>$\beta_2 = 100$ | N/A<br>N/A<br>N/A<br>N/A<br>N/A<br>N/A | 1 | 1 |
| | $\mu = 1,451.220664$<br>$\theta = 20$<br>$k = 19.986089$<br>$\gamma = 1,451.220664$<br>$\beta_1 = 100$<br>$\beta_2 = 100$ | N/A<br>N/A<br>N/A<br>N/A<br>N/A<br>N/A | 1 | 1 |
| | $\mu = 48,091.705355$<br>$\theta = 40$<br>$k = 15.184838$<br>$\gamma = 4,227.122626$<br>$\beta_1 = 100$<br>$\beta_2 = 100$ | N/A<br>N/A<br>N/A<br>N/A<br>N/A<br>N/A | 1 | 1 |

| | | | | |
|---|---|---|---|---|
| | $\mu = 17,386.031616$<br>$\theta = 5$<br>$k = 24.387861$<br>$\gamma = 4,722.311054$<br>$\beta_1 = 1,000$<br>$\beta_2 = 1,000$ | N/A<br>N/A<br>N/A<br>N/A<br>N/A<br>N/A | 1 | 1 |
| | $\mu = 42,100.456732$<br>$\theta = 10$<br>$k = 23.897082$<br>$\gamma = 4,720.616247$<br>$\beta_1 = 1,000$<br>$\beta_2 = 1,000$ | N/A<br>N/A<br>N/A<br>N/A<br>N/A<br>N/A | 1 | 1 |
| | $\mu = 1,451.220664$<br>$\theta = 20$<br>$k = 19.986089$<br>$\gamma = 1,451.220664$<br>$\beta_1 = 1,000$<br>$\beta_2 = 1,000$ | N/A<br>N/A<br>N/A<br>N/A<br>N/A<br>N/A | 1 | 1 |
| | $\mu = 48,091.705355$<br>$\theta = 40$<br>$k = 15.184838$<br>$\gamma = 4,227.122626$<br>$\beta_1 = 1,000$<br>$\beta_2 = 1,000$ | N/A<br>N/A<br>N/A<br>N/A<br>N/A<br>N/A | 1 | 1 |
| | $\mu = 100,000$<br>$\theta = 80$<br>$k = 10.478416$<br>$\gamma = 300$<br>$\beta_1 = 791.529945$<br>$\beta_2 = 791.529945$ | N/A<br>N/A<br>N/A<br>N/A<br>N/A<br>N/A | 1 | 1 |
| | $\mu : 2,651.085082205 \rightarrow 4,923.443724094999$<br>$\theta = 5$<br>$k : 10.90354305236 \rightarrow 20.249437097239998$<br>$\gamma : 1,753.786897891 \rightarrow 3,257.032810368999$<br>$\beta_1 = 100$<br>$\beta_2 = 100$ | uniform<br>N/A<br>uniform<br>uniform<br>N/A<br>N/A | 50 | 50 |
| | $\mu : 1,710.242215311 \rightarrow 3,176.164114148999$<br>$\theta = 10$<br>$k : 283.6877452059 \rightarrow 283.6877452059$<br>$\gamma : 1,094.775519327 \rightarrow 2,033.154535893$<br>$\beta_1 = 100$<br>$\beta_2 = 100$ | uniform<br>N/A<br>uniform<br>uniform<br>N/A<br>N/A | 50 | 50 |

| | | | | |
|---|---|---|---|---|
| | $\mu : 1,515.330622015 \rightarrow 2,814.185440885$<br>$\theta = 20$<br>$k : 45.097394656999995 \rightarrow 83.752304362999993$<br>$\gamma : 1,627.661264775 \rightarrow 3,022.799491724999$<br>$\beta_1 = 100$<br>$\beta_2 = 100$ | uniform<br>N/A<br>uniform<br>uniform<br>N/A<br>N/A | 50 | 50 |
| | $\mu : 3,358.179165312999 \rightarrow 6,236.618449866999$<br>$\theta = 40$<br>$k : 40.382682841689999 \rightarrow 74.996410991709993$<br>$\gamma : 3,200.381446073 \rightarrow 5,943.565542707$<br>$\beta_1 = 100$<br>$\beta_2 = 100$ | uniform<br>N/A<br>uniform<br>uniform<br>N/A<br>N/A | 50 | 50 |
| | $\mu : 2,110.31603188 \rightarrow 3,919.158344919999$<br>$\theta = 80$<br>$k : 18.47568732737 \rightarrow 34.311990750829999$<br>$\gamma : 2,464.694742762 \rightarrow 4,577.290236557999$<br>$\beta_1 = 100$<br>$\beta_2 = 100$ | uniform<br>N/A<br>uniform<br>uniform<br>N/A<br>N/A | 50 | 50 |
| | $\mu = 10,000$<br>$\theta = 10$<br>$k = 405.268207437$<br>$\gamma = 1,563.96502761$<br>$\beta_1 = 100$<br>$\beta_2 = 100$ | N/A<br>N/A<br>N/A<br>N/A<br>N/A<br>N/A | 1 | 1 |
| | $\mu = 10,000$<br>$\theta = 20$<br>$k = 64.42484951$<br>$\gamma = 2,325.23037825$<br>$\beta_1 = 100$<br>$\beta_2 = 100$ | N/A<br>N/A<br>N/A<br>N/A<br>N/A<br>N/A | 1 | 1 |
| | $\mu = 10,000$<br>$\theta = 40$<br>$k = 57.6895469167$<br>$\gamma = 4,571.97349439$<br>$\beta_1 = 100$<br>$\beta_2 = 100$ | N/A<br>N/A<br>N/A<br>N/A<br>N/A<br>N/A | 1 | 1 |

| | | | | |
|---|---|---|---|---|
| | $\mu = 10,000$<br>$\theta = 80$<br>$k = 26.3938390391$<br>$\gamma = 3,520.99248966$<br>$\beta_1 = 100$<br>$\beta_2 = 100$ | N/A<br>N/A<br>N/A<br>N/A<br>N/A<br>N/A | 1 | 1 |
| | $\mu = 4,280.2407818$<br>$\theta = 40$<br>$k = 77.947559070599993$<br>$\gamma = 4,850.84752348$<br>$\beta_1 = 100$<br>$\beta_2 = 100$ | N/A<br>N/A<br>N/A<br>N/A<br>N/A<br>N/A | 1 | 1 |
| | $\mu = 10,000$<br>$\theta = 40$<br>$k = 77.947559070599993$<br>$\gamma = 4,850.84752348$<br>$\beta_1 = 100$<br>$\beta_2 = 100$ | N/A<br>N/A<br>N/A<br>N/A<br>N/A<br>N/A | 1 | 1 |
| | $\mu = 4,280.2407818$<br>$\theta = 40$<br>$k = 77.947559070599993$<br>$\gamma = 10,000$<br>$\beta_1 = 100$<br>$\beta_2 = 100$ | N/A<br>N/A<br>N/A<br>N/A<br>N/A<br>N/A | 1 | 1 |
| | $\mu = 4,280.2407818$<br>$\theta = 40$<br>$k = 150$<br>$\gamma = 4,850.84752348$<br>$\beta_1 = 100$<br>$\beta_2 = 100$ | N/A<br>N/A<br>N/A<br>N/A<br>N/A<br>N/A | 1 | 1 |
| | $\mu = 327.213726$<br>$\theta = 1$<br>$k = 0.334031$<br>$\gamma = 100.686086$<br>$\beta_1 = 100$<br>$\beta_2 = 100$ | N/A<br>N/A<br>N/A<br>N/A<br>N/A<br>N/A | 5 | 5 |
| | $\mu = 281.863204$<br>$\theta = 1$<br>$k = 0.071928$<br>$\gamma = 102.204628$<br>$\beta_1 = 100$<br>$\beta_2 = 100$ | N/A<br>N/A<br>N/A<br>N/A<br>N/A<br>N/A | 5 | 5 |
| | $\mu = 1,730.059535$<br>$\theta = 5$<br>$k = 25.722558$<br>$\gamma = 3,170.20062$<br>$\beta_1 = 100$<br>$\beta_2 = 100$ | N/A<br>N/A<br>N/A<br>N/A<br>N/A<br>N/A | 5 | 5 |

| | | | | |
|---|---|---|---|---|
| | $\mu = 1,940.075003$<br>$\theta = 5$<br>$k = 27.569197$<br>$\gamma = 2,349.697838$<br>$\beta_1 = 100$<br>$\beta_2 = 100$ | N/A<br>N/A<br>N/A<br>N/A<br>N/A<br>N/A | 5 | 5 |
| | $\mu = 4,234.730823$<br>$\theta = 10$<br>$k = 23.233486$<br>$\gamma = 4,698.609761$<br>$\beta_1 = 100$<br>$\beta_2 = 100$ | N/A<br>N/A<br>N/A<br>N/A<br>N/A<br>N/A | 5 | 5 |
| | $\mu = 3,745.295176$<br>$\theta = 10$<br>$k = 20.974537$<br>$\gamma = 4,722.311054$<br>$\beta_1 = 100$<br>$\beta_2 = 100$ | N/A<br>N/A<br>N/A<br>N/A<br>N/A<br>N/A | 5 | 5 |
| | $\mu = 4,147.742155$<br>$\theta = 20$<br>$k = 19.026819$<br>$\gamma = 4,868.321675$<br>$\beta_1 = 100$<br>$\beta_2 = 100$ | N/A<br>N/A<br>N/A<br>N/A<br>N/A<br>N/A | 5 | 5 |
| | $\mu = 4,627.282885$<br>$\theta = 20$<br>$k = 16.868596$<br>$\gamma = 1,273.002093$<br>$\beta_1 = 100$<br>$\beta_2 = 100$ | N/A<br>N/A<br>N/A<br>N/A<br>N/A<br>N/A | 5 | 5 |
| | $\mu = 48,091.705355$<br>$\theta = 40$<br>$k = 15.184838$<br>$\gamma = 4,227.122626$<br>$\beta_1 = 100$<br>$\beta_2 = 100$ | N/A<br>N/A<br>N/A<br>N/A<br>N/A<br>N/A | 5 | 5 |
| | $\mu = 10,000$<br>$\theta = 80$<br>$k = 9.721245$<br>$\gamma = 300$<br>$\beta_1 = 79.152995$<br>$\beta_2 = 79.152995$ | N/A<br>N/A<br>N/A<br>N/A<br>N/A<br>N/A | 5 | 5 |
| | $\mu = 48,091.705355$<br>$\theta = 40$<br>$k = 15.184838$<br>$\gamma = 4,227.122626$<br>$\beta_1 = 150$<br>$\beta_2 = 150$ | N/A<br>N/A<br>N/A<br>N/A<br>N/A<br>N/A | 3 | 3 |

| | | | | |
|---|---|---|---|---|
| | $\mu = 72,137.5580325$<br>$\theta = 40$<br>$k = 15.184838$<br>$\gamma = 4,227.122626$<br>$\beta_1 = 150$<br>$\beta_2 = 150$ | N/A<br>N/A<br>N/A<br>N/A<br>N/A<br>N/A | 3 | 3 |
| | $\mu = 100,000$<br>$\theta = 40$<br>$k = 15.184838$<br>$\gamma = 4,227.122626$<br>$\beta_1 = 150$<br>$\beta_2 = 150$ | N/A<br>N/A<br>N/A<br>N/A<br>N/A<br>N/A | 3 | 3 |
| | $\mu\ :\ 77,000 \to 140,000$<br>$\theta = 40$<br>$k\ :\ 1.5 \to 100.5$<br>$\gamma\ :\ 3,200 \to 5,000$<br>$\beta_1\ :\ 75 \to 165$<br>$\beta_2 = \beta_1$ | uniform<br>N/A<br>uniform<br>uniform<br>uniform<br>N/A | 100 | 58 |
| | $\mu\ :\ 7,700 \to 14,000$<br>$\theta = 20$<br>$k\ :\ 1.5 \to 100.5$<br>$\gamma\ :\ 3,200 \to 5,000$<br>$\beta_1\ :\ 75 \to 165$<br>$\beta_2 = \beta_1$ | uniform<br>N/A<br>uniform<br>uniform<br>uniform<br>N/A | 18 | 13 |
| | $\mu\ :\ 4,600 \to 10,000$<br>$\theta = 40$<br>$k\ :\ 1.5 \to 100.5$<br>$\gamma\ :\ 1,200 \to 3,000$<br>$\beta_1\ :\ 75 \to 165$<br>$\beta_2 = \beta_1$ | uniform<br>N/A<br>uniform<br>uniform<br>uniform<br>N/A | 308 | 230 |
| | $\mu\ :\ 2,000 \to 5,600$<br>$\theta = 20$<br>$k\ :\ 20 \to 416$<br>$\gamma\ :\ 140 \to 500$<br>$\beta_1 = 100$<br>$\beta_2 = \beta_1$ | uniform<br>N/A<br>uniform<br>uniform<br>N/A<br>N/A | 100 | 68 |
| Idealized sensor-actuated AIF | $\mu\ :\ 0.01\theta \to 100\theta$<br>$\theta\ :\ 0.01 \to 100$<br>$\gamma\ :\ 0.01 \to 100$<br>$\nu\ :\ 1.1\mu\beta_x/\theta \to 1.5\mu\beta_x/\theta$<br>$\kappa = 0.05$<br>$n = 1 - \sqrt{1 + \theta/\beta_x}$ | logarithmic<br>logarithmic<br>logarithmic<br>uniform<br>N/A<br>N/A | $5,000$ | $3,985$ |
| | $\mu\ :\ 0.01\theta \to 100\theta$<br>$\theta\ :\ 0.01 \to 100$<br>$\gamma\ :\ 0.01 \to 100$<br>$\nu\ :\ 1.1\mu\beta_x/\theta \to 1.5\mu\beta_x/\theta$<br>$\kappa = 0.05$<br>$n\ :\ 0.1\alpha \to 10\alpha$ | logarithmic<br>logarithmic<br>logarithmic<br>uniform<br>N/A<br>uniform | $5,000$ | $3,943$ |

| | | | | |
|---|---|---|---|---|
| | $\alpha \coloneqq 1 - \sqrt{1+\theta/\beta_x}$ | N/A | | |
| | $\mu$ : $0.01\theta \to 100\theta$<br>$\theta$ : $0.01 \to 100$<br>$\gamma$ : $0.05 \to 50$<br>$\nu$ : $1.1\mu\beta_x/\theta \to 10\mu\beta_x/\theta$<br>$\kappa = 1$<br>$n$ : $0.1\alpha \to 10\alpha$<br>$\alpha \coloneqq 1 - \sqrt{1+\theta/\beta_x}$ | logarithmic<br>logarithmic<br>logarithmic<br>uniform<br>N/A<br>uniform<br>N/A | $7,500$ | $4,032$ |
| | $\mu$ : $5 \to 95$<br>$\theta$ : $5 \to 95$<br>$\gamma$ : $5 \to 95$<br>$\nu$ : $1.1\mu\beta_x/\theta \to 10\mu\beta_x/\theta$<br>$\kappa = 1$<br>$n$ : $0.1\alpha \to 10\alpha$<br>$\alpha \coloneqq 1 - \sqrt{1+\theta/\beta_x}$ | uniform<br>uniform<br>uniform<br>uniform<br>N/A<br>uniform<br>N/A | $7,495$ | $7,376$ |
| | $\mu$ : $0.1\theta \to 100\theta$<br>$\theta$ : $5 \to 95$<br>$\gamma$ : $0.1 \to 100$<br>$\nu$ : $1.1\mu\beta_x/\theta \to 10\mu\beta_x/\theta$<br>$\kappa = 1$<br>$n$ : $0.1\alpha \to 10\alpha$<br>$\alpha \coloneqq 1 - \sqrt{1+\theta/\beta_x}$ | logarithmic<br>uniform<br>logarithmic<br>uniform<br>N/A<br>uniform<br>N/A | $4,952$ | $4,916$ |
| | $\mu$ : $0.1\theta \to 100\theta$<br>$\theta$ : $20 \to 100$<br>$\gamma$ : $0.1 \to 100$<br>$\nu$ : $1.1\mu\beta_x/\theta \to 100\mu\beta_x/\theta$<br>$\kappa$ : $0.1 \to 1$<br>$n$ : $0.5\alpha \to 2\alpha$<br>$\alpha \coloneqq 1 - \sqrt{1+\theta/\beta_x}$ | logarithmic<br>uniform<br>logarithmic<br>uniform<br>N/A<br>uniform<br>N/A | $10,000$ | $3,268$ |
